# Abnormal activation of the mineralocorticoid receptor in the aldosterone-sensitive distal nephron contributes to fructose-induced salt-sensitive hypertension

**DOI:** 10.1101/2024.08.19.608663

**Authors:** Ronghao Zhang, Shujie Shi, Darshan Aatmaram Jadhav, Najeong Kim, Autumn Brostek, Beau R. Forester, Rashi Shukla, Christina Qu, Benjamin Kramer, Jeffrey L. Garvin, Thomas R. Kleyman, Agustin Gonzalez-Vicente

## Abstract

Fructose high-salt (FHS) diets increase blood pressure (BP) in an angiotensin II (Ang II)-dependent manner. Ang II stimulates aldosterone release, which, by acting on the mineralocorticoid receptor (MR), regulates Na^+^ reabsorption by the aldosterone-sensitive distal nephron (ASDN). The MR can be transactivated by glucocorticoids, including those locally produced by 11β-HSD1. The epithelial sodium channel (ENaC) is a key transporter regulated by MRs. We hypothesized that fructose-induced salt-sensitive hypertension depends in part on abnormal activation of MRs in the ASDN with consequent increases in ENaC expression. We found that aldosterone-upregulated genes in mice ASDN, significantly overlapped with 74 genes upregulated by FHS in the rat kidney cortex (13/74; p≤1x10^-8^), and that these 74 genes are prominently expressed in rat ASDN cells. Additionally, the average z-score expression of mice-aldosterone-upregulated genes is highly correlated with FHS compared to glucose high-salt (GHS) in the rat kidney cortex (Pearson correlation; r=0.66; p≤0.005). There were no significant differences in plasma aldosterone concentrations between the FHS and GHS. However, 11β-HSD1 transcripts were upregulated by FHS (log_2_FC=0.26, p≤0.02). FHS increased BP by 23±6 mmHg compared to GHS, and blocking MRs with eplerenone prevented this increase. Additionally, inhibiting ENaC with amiloride significantly reduced BP in FHS from 148±6 to 134±5 mmHg (p≤0.019). Compared to GHS, FHS increased total and cleaved αENaC protein by 89±14 % (p≤0.03) and 47±16 % (p≤0.01) respectively. FHS did not change β- or γ-subunit expression. These results suggest that fructose-induced salt-sensitive hypertension depends, in part, on abnormal Na^+^ retention by ENaC, resulting from the activation of MRs by glucocorticoids.

## Introduction

Hypertension is now the leading cause of “loss of health” worldwide (1). Hypertension increases the likelihood of stroke, vascular disease, kidney failure, heart failure, and blindness. Nearly one in two people in the United States (US) has high blood pressure (BP) (2). In salt-sensitive hypertension, BP increases with salt consumption. This disease may account for up to 50% of US hypertension (3), and is particularly prevalent in African Americans (3-7). Salt-sensitive hypertension must include a renal defect or pressure natriuresis would restore normal BP (8, 9).

Fructose consumption in the US has increased dramatically since 1970. The diet of the average American now contains more than 11% of its calories as fructose while >17 million Americans ingest >20% of their calories as fructose (10-12). The increase in fructose consumption mirrors the rise in the incidence of hypertension in the US from 18% to >40% suggesting a link between the two. In fact, several studies have shown that fructose causes hypertension (13-15), including salt-sensitive hypertension (16-18). Fructose in an amount that mimics a single soft drink increased both mean and diastolic BP in healthy adults more than glucose or sucrose in a randomized, crossover study (19). Large cross-sectional studies show that dietary fructose is linked with hypertension (14, 15, 20). Even the average amount of fructose consumed by the average American (11% of calories) likely contributes to both elevated BP and salt sensitivity (21). However, the mechanisms by which dietary fructose induces salt-sensitive hypertension remain unclear.

In animals, fructose-induced salt-sensitive hypertension is angiotensin II (Ang II)-dependent. Losartan, an Ang II type I receptor antagonist, prevents the increase in BP (22). Ang II regulates BP via pleiotropic actions (23-25). In addition to its vascular effects (25), Ang II stimulates Na^+^ reabsorption in the proximal tubule (PT) (26-28). Studies from our lab have shown that FHS increases the sensitivity to Ang II in the proximal nephron, shifting the concentration-response curve for Na^+^ reabsorption (29, 30) and O_2_^-^ production (22, 31, 32) towards lower concentrations (left shift). Ang II also controls the release of aldosterone (33), a hormone that augments Na^+^ reabsorption by the late connecting tubule and collecting duct (34-37); commonly known as the aldosterone-sensitive distal nephron (ASDN). Aldosterone exerts its effects by acting on the mineralocorticoid receptor (MR), which can also be activated by the glucocorticoid corticosterone. To prevent the inappropriate activation of the MR by corticosterone, the aldosterone-sensitive tissues express hydroxysteroid 11-beta dehydrogenase 2 (11β-HSD2; *Hsd11b2*), which converts corticosterone to its inactive form, cortisone. However, the actions of 11β-HSD2 can be counteracted by hydroxysteroid 11-beta dehydrogenase 1 (11β-HSD1; *Hsd11b1*). This enzyme converts cortisone to corticosterone, allowing tissue-specific regulation of corticosterone levels and paracrine MR activation. Upon activation, the MR migrates to the nucleus and initiates the transcription of genes involved in Na^+^ transport. One of its primary targets is the epithelial sodium channel (ENaC), expressed throughout the ASDN (38). ENaC is composed of three subunits: alpha, beta, and gamma, which are in part regulated by proteolytic cleavage (39, 40). Previous studies by us (31) and others (41) suggest that in addition to increased transport by proximal tubules, fructose may also affect the ASDN. However, whether aldosterone and/or the MR play a relevant role in fructose-induced salt-sensitive hypertension has not been studied. We hypothesized that fructose-induced salt-sensitive hypertension depends in part on abnormal activation of MRs in the ASDN with consequent increases in ENaC expression.

## Methods

### Animals

All animal studies were approved by the Case Western Reserve University Institutional Animal Care and Use Committee following the National Institutes of Health Guidelines for Use and Care of Laboratory Animals. Wild-type Sprague-Dawley rats were purchased from Charles River Breeding Laboratories (Wilmington, MA; Raleigh, NC; Kingston, NY). All experiments were conducted in male animals, unless noted in the protocols.

### Dietary intervention

Upon arrival at our animal facility, rats were fed a standard rodent diet (Prolab® IsoPro® RMH 3000) containing normal salt (NS; 0.26% Na^+^). Animals were allowed to acclimate a few days before being randomly assigned to experimental diet groups. The experimental solid diets were previously used by us (22, 31) and contained 4.0% NaCl, and either 20% glucose (GHS; Test Diet #9GSH) or 20% fructose (FHS; Test Diet #9GSK). The other components of both diets were the same: 1.5% Na^+^, 1.2% K^+^, 10% Fat, 18.6% protein, 4.6% fiber, and 55.5% total carbohydrates. Animals have unrestricted access to food and water at all times.

### Bulk kidney cortex transcriptomes

We used kidney cortex RNA sequencing (RNAseq) transcriptomes from Groups 7 and 8 in our previous work, Forester et.al 2024 (42). These datasets consisted of 6 FHS transcriptomes (Group 7), and 6 GHS transcriptomes (Group 8) from wild-type Sprague-Dawley male rats. RNA was extracted using standard methods (42, 43), and RNAseq data pre-processing, quality control, alignment, and count normalization were conducted using previously reported pipelines (42-44). Raw sequencing files were made available at xxxTBD.

### Network analysis

We used the network produced in our previous publication (42). The gene coexpression network was constructed with the R package WGCNA: Weighted Correlation Network Analysis (v1.71) (45, 46) using a standard “Step by Step Method” workflow. For each gene, WGCNA calculated a Pearson correlation coefficient (r) along with its corresponding p-value, linking the gene’s expression level to fructose consumption. Here, FHS was used as the delta level and GHS as the baseline. A positive correlation signifies that the gene’s expression increases with fructose consumption, while a negative correlation indicates a decrease.

### Single-nuclei (sn) suspensions

We produced a snRNAseq dataset from two different pools of kidney cortexes: one from 3 male and one from 3 female wild-type Sprague Dawley rats, all fed standard rodent diet (NS, Prolab® IsoPro® RMH 3000). Single nuclei suspensions were obtained using the protocol published by the Kidney Precision Medicine Project (KPMP) (47), with minor adaptations. In brief, ∼5 mg of kidney cortex collected from anesthetized rats were homogenized in a Dounce homogenizer. Lysis buffer contained: 20 mmol/l Tris (pH 8.0), 320 mmol/l sucrose, 5 mmol/l CaCl_2_, 3 mmol/l Mg(CH_3_COO)_2_, 0.1 mmol/l EDTA, 100 mmol/l Triton X-100, 5 µg/ml DAPI and 400 U/ml RNAse inhibitors cocktail (Qiagen, Y9240L) dissolved in molecular biology grade water. Nuclei in suspension were passed through a 30 µm filter and washed. After washing, nuclei were spun by centrifugation (500 xg, 10 min, 4°C), and supernatant discarded. Nuclei were resuspended in adequate volume of PBS added 400 U/ml of RNAse inhibitors to a concentration of ∼5x10^5^ nuclei/ml. Nuclei concentration was measured by DAPI fluorescence using Countess™ 3 Automated Cell Counter (Invitrogen, AMQAX2000).

### Library construction

Libraries were constructed in our lab using Chromium GEM-X Single Cell 3’ Reagent Kit v4 according to manufacturers’ recommendations. In brief, appropriate nuclei suspension volume was added to the master mix to target a recovery of 20,000 nuclei. Samples were loaded onto a GEM-X 3’ Chip to generate the emulsion. Reverse transcription was conducted in a PTC Tempo (Bio-Rad, #12015392) thermal cycler with a standard protocol. The resulting 10X barcoded full-length cDNA was cleaned with Dynabeads MyOne SILANE, and quality control was conducted with an Agilent 5200 Fragment Analyzer prior to amplification. Amplified cDNA was cleaned and quality controlled. Finally, the 3’ gene expression library was constructed with GEX fragmentation, ligation, and final cleanup.

### Sequencing and processing of snRNA transcriptomes

The Genomics Core at Case Western Reserve University (CWRU) School of Medicine sequenced the snRNA library on a NovaSeq X system. Raw sequencing files were made available at xxxTBD. Data was processed in our servers using methods similar to those we described before (44). In brief, count matrices were obtained on CellRanger (v8.0), using the rat mRatBN7.2 genome as the reference. Ambient RNA was removed using SoupX (v1.6.2.). Quality control was conducted using Seurat (v5.1.0), keeping features detected in at least three cells. The following filters were applied: nFeature_RNA > 400 and percent.MT < 20. The QC process yielded a merged dataset consisting of 52,206 cells and 23,882 features. The dataset was log-normalized, and cell-cycle scores were calculated using the updated 2019 cc.genes, which were also regressed out during data scaling. Potential doublets were identified and eliminated using DoubletFinder (v2.0.4) with parameters inferred via *paramSweep()*. *SCTransform()* was applied to male and female pool datasets individually, and then both were integrated using *IntegrateData()* with 3,000 integration features. Dimensionality reduction was conducted, and 44 clusters were identified at a resolution of 3.6, employing the Louvain algorithm with *FindClusters()*. For assigning clusters to specific cell types, we used the snRNAseq map (v1.5) from the KPMP atlas as the reference. Ortholog genes were identified using Ensembl’s BioMart, and both datasets were harmonized after similar normalization, scaling, and dimensionality reduction. The *FindTransferAnchors()* function was then employed. *TransferData()* was used to transfer cell type information, for subclass.l2 (cell type), and then cortical epithelial cell types were extracted, while label assignments for medullary epithelial cells and classes different than epithelial were discarded.

### Module Scores

Using bulk kidney cortex transcriptomes, we previously identified a WGCNA module containing cell annotations from the ASDN (31). To study whether this module, named paleturquoise, was enriched with genes expressed in the ASDN, we calculated module scores with the *AddModuleScore()* function from Seurat (v5.1.0) using the newly generated snRNAseq data. Transferred labels of cortical epithelial cells from subclass.l2 were assigned to different sections of the nephron as follows: podocytes and parietal epithelial cells were assigned to the “glomerular epithelial cell” (GEC), PT segments S1 and S2 were assigned to the “proximal tubule”, cortical thick ascending limb and macula densa cells were assigned to the “loop of Henle” (LOH), and distal convoluted tubule cell subtypes 1 and 2, connecting tubule cells, and cortical collecting duct principal and intercalating cells were assigned to the ASDN.

### Distal nephron aldosterone signature

An aldosterone-induced genomic signature on the ASDN was obtained by identifying genes whose expression changes in response to aldosterone on murine ASDN epithelial cells (37). The dataset contained transcriptomes of late distal convoluted tubule cells (DCT2) connecting tubule (CNT) cells and the initial cortical collecting duct (iCCD) principal cells from mice treated with vehicle or aldosterone for 6 days. Differential expression data were downloaded from the National Heart, Lung, and Blood Institute (NHLBI-NIH) Epithelial Systems Biology Laboratory (48). The mouse transcriptomes were first mapped to our rat kidney cortex transcriptomes with a shared space of 8618 genes. Within the shared gene space, aldosterone-response genes were identified as those with a p-value ≤ 0.05, and either log_2_Ratio ≥ 0.5 (Upregulated) or ≤ -0.5 (Downregulated). We identified 203 upregulated and 214 downregulated genes by aldosterone, while 8201 did not present significant changes.

### Mapping the mice ASDN aldosterone signature to WGCNA modules

We used the R function *fisher.test()* with the parameter ‘alternative = “greater”’ to identify positive enrichments between WGCNA modules and the murine aldosterone signature from the ASDN. The shared gene space (8201 genes) was used as the background list. We found that only the 203 aldosterone-upregulated genes presented significant overlaps with WGCNA modules, while the 214 downregulated genes did not. Thus, we only focused on the upregulated gene space.

### Aldosterone response genes in fructose

Only genes upregulated by aldosterone the mouse ASDN significantly overlap with WGCNA modules. Thus, we studied the expression of those genes in response to FHS in the rat cortex. We calculated genomewide z-scores for each rat, 6 FHS (Group 7, Forester et.al 2024 (42)), and 6 GHS (Group 8, Forester et.al 2024 (42)). Then we extracted the 203 genes upregulated by aldosterone in the mouse ASDN and compared the averaged z-score of each rat.

### Plasma aldosterone concentrations

Aldosterone was measured in fresh plasma of rats fed either NS, GHS or FHS using the Aldosterone Competitive ELISA Kit (Invitrogen, Cat. EIAALD) according to the manufacturer’s recommendations. In brief, 250µl of serum was extracted with ethyl-acetate and dried in a vacuum concentrator (ThermoFisher, #DNA130). The resulting pellet was reconstituted in 10µl of ethanol and taken to 250µl with the provided buffer before running the competitive assay. Competitive binding data were analyzed with the R package drc (drc_3.0-1). The calibration data were fitted to a four-parameter log-logistic model using *drm(*Optical Density ∼ Standard Concentration), estimating the parameters “Slope”, “Lower”, “Upper” and “Km”, corresponding to slope, lower limit, upper limit, and the concentration at half-maximal binding, respectively. Finally, the *ED()* function was used to determine the aldosterone concentrations corresponding to the optical densities of the samples.

### Differential gene expression

Genomewide differential gene expression on bulk kidney cortex transcriptomes was performed using the R package DESeq2 (v1.34.0.) similar to our previous report (42). The formula (design = ∼ dietary treatment) was used to identify changes in gene expression caused by the FHS (Group 7, n=6) as compared to GHS (Group 8, n=6), used as the base level. Differentially expressed genes were defined by the following criteria: 1) Consistent directionality between WGCNA (Pearson correlation) and DESeq2 (Wald test) analyses; 2) A significant Pearson correlation coefficient with FHS (-0.5 ≤ r ≥ 0.5; p ≤ 0.05) and 3) An absolute log_2_ fold change (log_2_FC) ≥ 0.25 (-0.25 ≤ log_2_FC ≥ 0.25; p ≤ 0.05) in the Wald test. A comprehensive table presenting the results from both analyses is provided as **Supplemental Table 1**.

### Blood pressure by plethysmography

Systolic BP was recorded by tail plethysmography using a CODA Non-invasive BP System (Kent Scientific Corporation, Torrington, CT) in male rats as we did before (22, 30, 31, 49). Briefly, rats were trained 2 or 3 times the week before the start of the dietary intervention while on the standard rodent diet (NS; Prolab® IsoPro® RMH 3000). Training consisted of restraining the animals, placing them on a temperature-controlled pad, positioning the cuffs and inflating them. Training BPs were not recorded. During training, rats were randomly assigned to one of three experimental groups: GHS (n=6), FHS (n=6) or FHS+E, given eplerenone in the drinking water to a resulting dose of ∼100 mg/kg body weight. BP training continued during the dietary intervention, and systolic BP was measure between days 7 and 8. The FHS+E group was euthanized after day 7-8. On day 9, BP was measured in the GHS and FHS animals, and then amiloride (A) was added to their drinking water at a concentration of 10^-4^ mole/l resulting in a dose of ∼4 mg/kg/day. On day 10 the BP from groups GHS+A and FHS+A was measured again.

### Western Blotting

Rats were anesthetized by intraperitoneal injection of 100 mg/kg ketamine and 20 mg/kg xylazine. An incision was made to expose the left kidney, and it was cooled *in situ* with 150 mM ice-cold NaCl. After 1-2 min, kidneys were excised and plunged into liquid nitrogen for 5 min. Flash-frozen kidney cortex pieces were collected by fracturing kidneys and stored at -80°C for long-term storage. Samples were homogenized in CelLytic™ MT lysis buffer (Sigma, #C3228) with added protease (1:100, Sigma, #539134) and phosphatase (1:100, Sigma, #P0044) inhibitors. Protein concentration was measured using a Pierce BCA protein assay kit (ThermoFisher, #23227). Lysates containing 50 μg of protein were incubated with an equal volume of 2X Laemmli buffer (Bio-Rad, #1610737, containing 10% β-mercaptoethanol) at 37 °C for 30 minutes. For blotting αENaC, samples were run on 10% polyacrylamide stain-free gels (Bio-Rad, #5678034), while for blotting βENaC and γENaC, 4-15% polyacrylamide gels were used (Bio-Rad, #5678084). Total protein per lane was detected with Bio-Rad Stain-Free® technology and used for normalization. Proteins were then transferred to nitrocellulose membranes (Millipore, #HAWP04700) and incubated with antibodies as shown in **Table 1**. To remove N-linked glycans from γENaC, a separate set of lysates was treated with Peptide-N-Glycosidase F (PNGase F; New England BioLabs, #P0704) according to the manufacturer’s recommendations and blotted as shown in **Table 1**. All blots were developed using Clarity Western ECL Substrates (Bio-Rad, #1705060) and imaged with a Bio-Rad ChemiDoc™ system. The intensity of the bands of interest was quantified using Bio-Rad Image Lab 6.1 and normalized to total protein loading in the individual lanes.

**Table 1:**
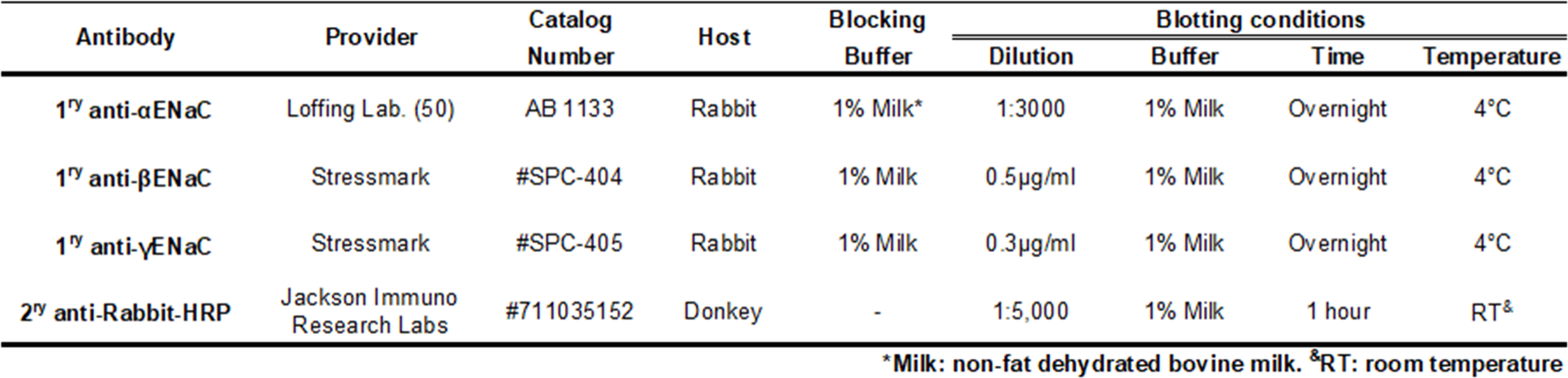
Antibodies and blotting conditions.

### Statistical Analysis

Unless reported in specific bioinformatics analyses statistical analyses were conducted using GraphPad Prism. Means and standard errors of the means (SEM) were calculated. Data were analyzed by one-way Analysis of Variance (ANOVA) or Student’s t-tests, paired or unpaired, as appropriate. Corrections for multiple comparisons were made using the Benjamini and Hochberg method. Nominal (p) or adjusted (p-Adj) p-values were considered significant when less than 0.05. Only significant p values are reported on the figures.

## Results

We based our hypothesis on our prior research (31) that identified genes whose expression changes in response to dietary fructose. Many of these genes were associated with ASDN-related enrichment terms, including “Aldosterone-regulated sodium reabsorption”. Thus, we studied the role of aldosterone and MR activation in fructose-induced salt-sensitive hypertension. First, using an aldosterone-transcriptional signature from the ASDN, we found that genes upregulated by aldosterone showed significant enrichment in three coexpression modules: paleturquoise (13 genes, p ≤ 1x10^-8^), darkred (17 genes, p ≤ 1x10^-7^) and sienna3 (24 genes, p ≤ 5x10^-28^) (**Figure 1-A**). Of those 3 modules, we focused downstream analyses on the paleturquoise, as it was associated with FHS in our previous work (31). Secondly, we used snRNAseq transcriptomes from the rat kidney cortex to generate module scores, showing that genes in the paleturquoise module are more highly expressed in the ASDN than in proximal tubules, thick ascending limbs, or the glomerular epithelium (**Figure 1-B**). These findings suggest that genes normally regulated by aldosterone in the ASDN are also affected by FHS.

**Figure 1:**
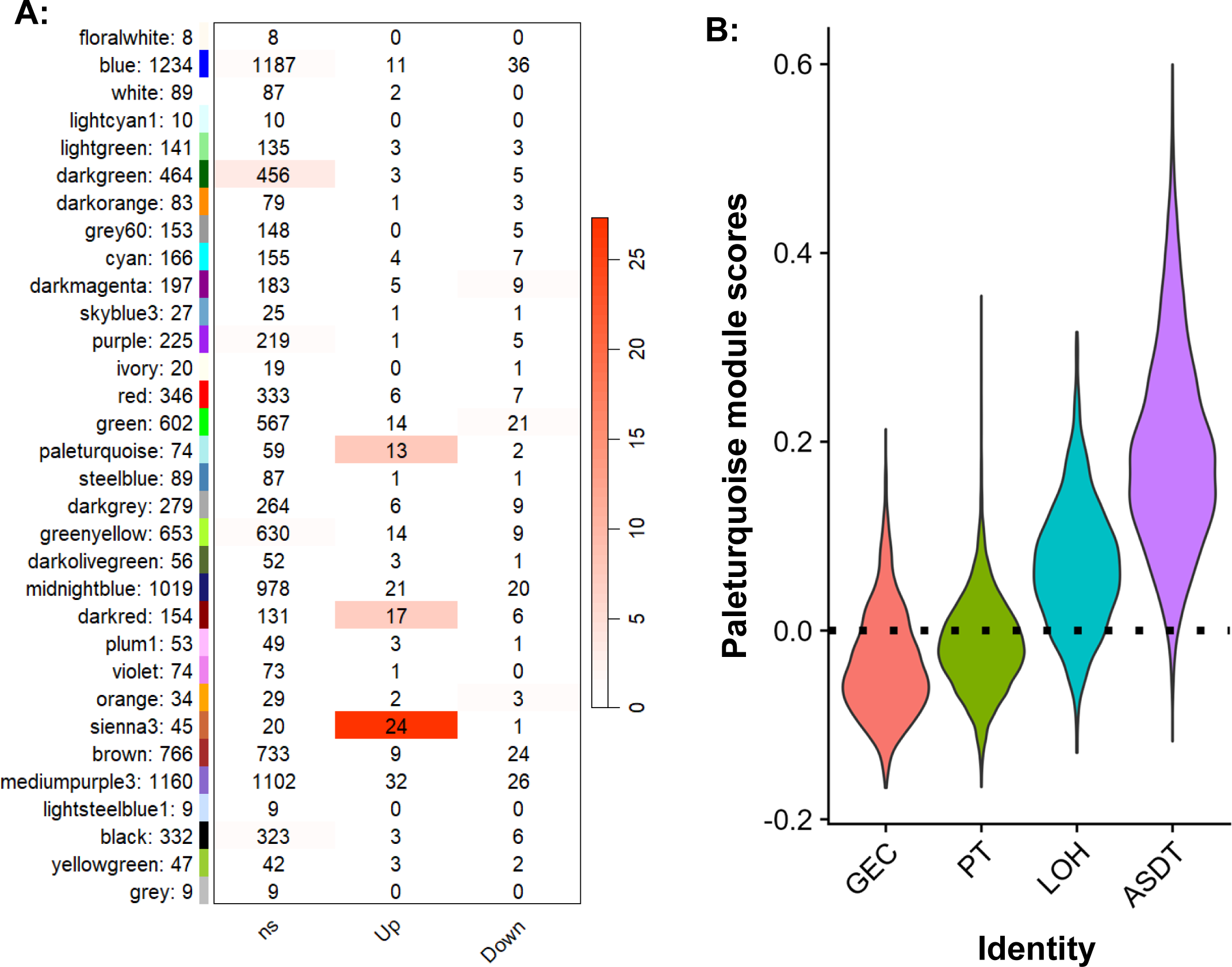
**A)** Gene overlaps between WGCNA modules and an aldosterone-induced signature in the mice distal nephron. The Fisher exact test showed that only three WGCNA modules presented significant overlaps with the genes upregulated by aldosterone, while there was no overlap on the downregulated ones. WGCNA modules and the number of in-module genes are shown on the left. The numbers on the table correspond to the intersecting genes. For the definition of upregulated (Up) or downregulated (Down) genes, refer to the main text, ‘ns’ means not significant. The color scale represents the log_10_(1/Fisher p-value). **B)** Module scores for the WGCNA paleturquoise module. The score represents the average expression level of genes within the paleturquoise module across different kidney cortex epithelial cell clusters of single-nuclei RNA transcriptomes from control animals. Genes in the paleturquoise module are most highly expressed in the aldosterone-sensitive distal nephron (ASDN) as compared to the proximal tubule (PT) Loop of Henle (LOH) or the glomerular epithelium (GEC).

The data presented above suggest increased activity of the MR in rats fed FHS. To support this hypothesis, we assess the downstream signaling of the MR in FHS and GHS animals by calculating the average z-score of the 203 aldosterone-upregulated genes in mice. We found a significant positive correlation coefficient (r = 0.75, p ≤ 0.005) between z-scores of genes upregulated by aldosterone and FHS (**Figure 2**). Then, we measured plasma aldosterone concentration. The excess dietary Na^+^ effectively suppressed plasma aldosterone levels from 187 ± 51 pg/ml (n = 5) in NS, to 56 ± 6 pg/ml in GHS (Δ = -131 ± 38 pg/ml; p-Adj ≤ 0.004; n = 6) and to 57 ± 6 pg/ml in FHS (Δ = -130 ± 38 pg/ml; p-Adj ≤ 0.004; n = 6) (**Figure 3**). There were no significant differences between GHS and FHS. Thus, plasma aldosterone levels cannot explain an increased activation of the MR in FHS.

**Figure 2:**
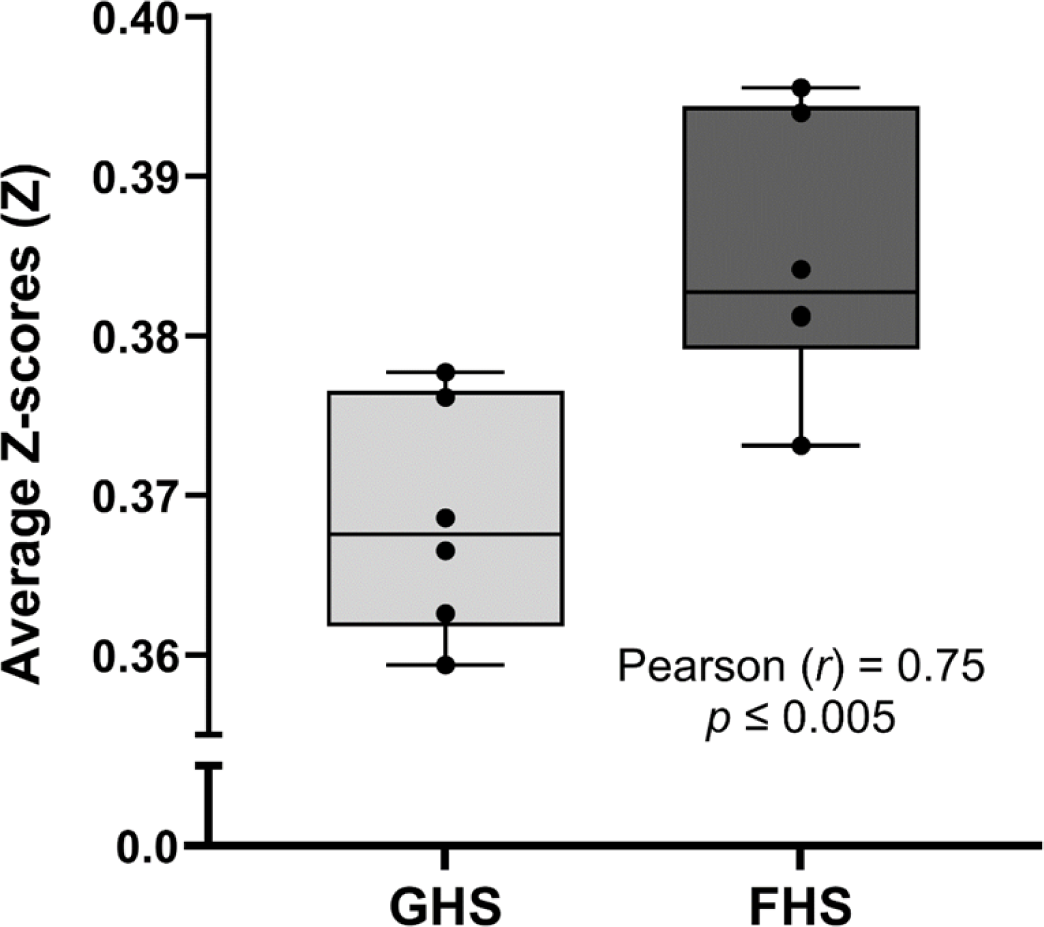
Average genomewide z-scores, a surrogate of gene activation in rats, of the 203 genes upregulated by aldosterone in the mice ASDN. Pearson correlation shows a significant association between the expressions of aldosterone-response genes and FHS (r = 0.75; p ≤ 0.005, n = 6).

**Figure 3:**
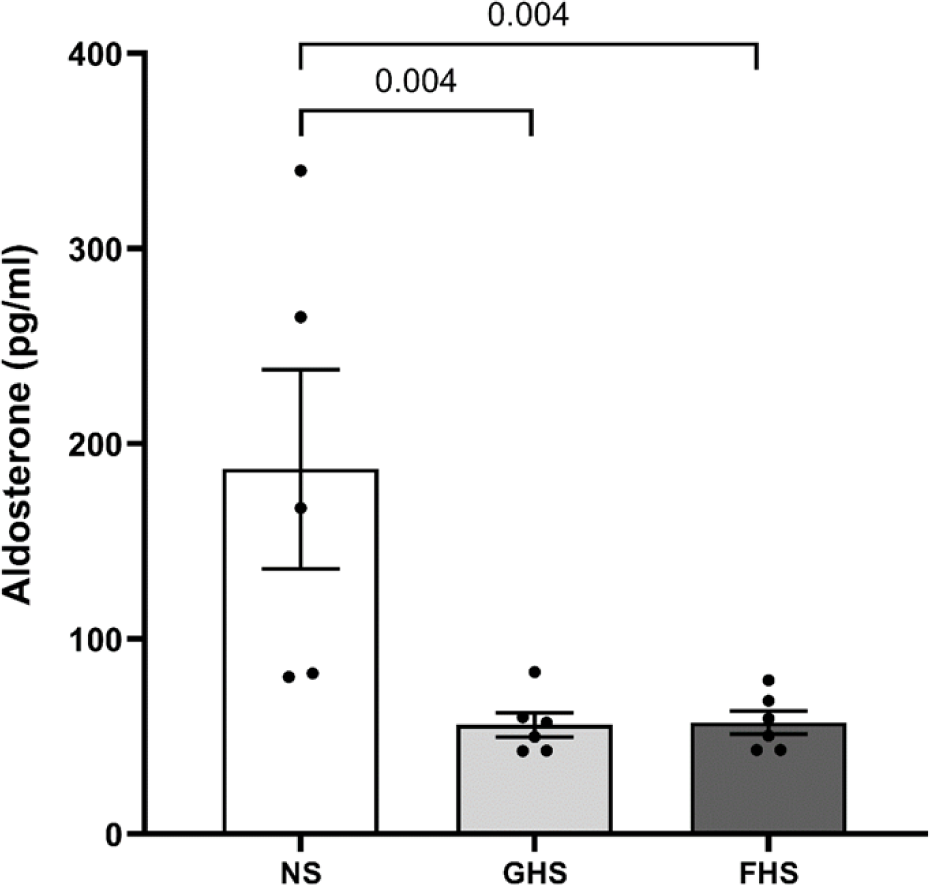
Aldosterone concentrations in plasma from rats fed regular chow containing normal salt (NS), glucose high-salt (GHS), or fructose high-salt (FHS). One-way ANOVA indicated that the means between groups were significantly different (p ≤ 0.006). Excess dietary Na^+^ effectively suppressed plasma aldosterone levels in both FHS and GHS. Plasma aldosterone presented no significant differences between FHS and GHS. Corrections for multiple comparisons were made using the Benjamini and Hochberg method. Adjusted p-values (p-Adj) are reported.

Tissues expressing the MR also express 11β-HSD2 to prevent activation by corticosterone. However, 11β-HSD1 converts cortisone to corticosterone, allowing local MR activation. We investigated differences in the 11β-HSD1 gene (*Hsd11b1*) expression in the kidney cortex between FHS and GHS in two ways: 1) using the Pearson correlation coefficient and significance between *Hsd11b1* and FHS, and 2) conducting differential gene expression with the Wald test. Our results show that *Hsd11b1* is positively correlated with the FHS diet (r ≥ 0.66; p ≤ 0.02) and the FHS diet significantly increases the expression of *Hsd11b1* (log_2_FC = 0.26, p ≤ 0.02). Together, these results suggest increased local levels of corticosterone in the kidney cortex of FHS.

To test whether MR activation contributes to fructose-induced salt-sensitive hypertension, we studied the effect of the MR blocker eplerenone on systolic BP in FHS-fed rats. We compared the final systolic BP of 3 groups of rats: FHS, FHS+E, and GHS. One-way ANOVA indicated that the means between groups were significantly different (p≤0.017). Post-test comparisons show that FHS significantly increased systolic BP as compared to GHS (p-Adj ≤ 0.044) within a week (**Figure 4**). Coadministration of eplerenone prevented the increase in systolic BP caused by FHS (p-Adj ≤ 0.0013). There were no differences in systolic BP between GHS and FHS+E (p-Adj ≤ 0.39). Together, these data indicate that increased MR signaling contributes to fructose-induced salt-sensitive hypertension.

**Figure 4:**
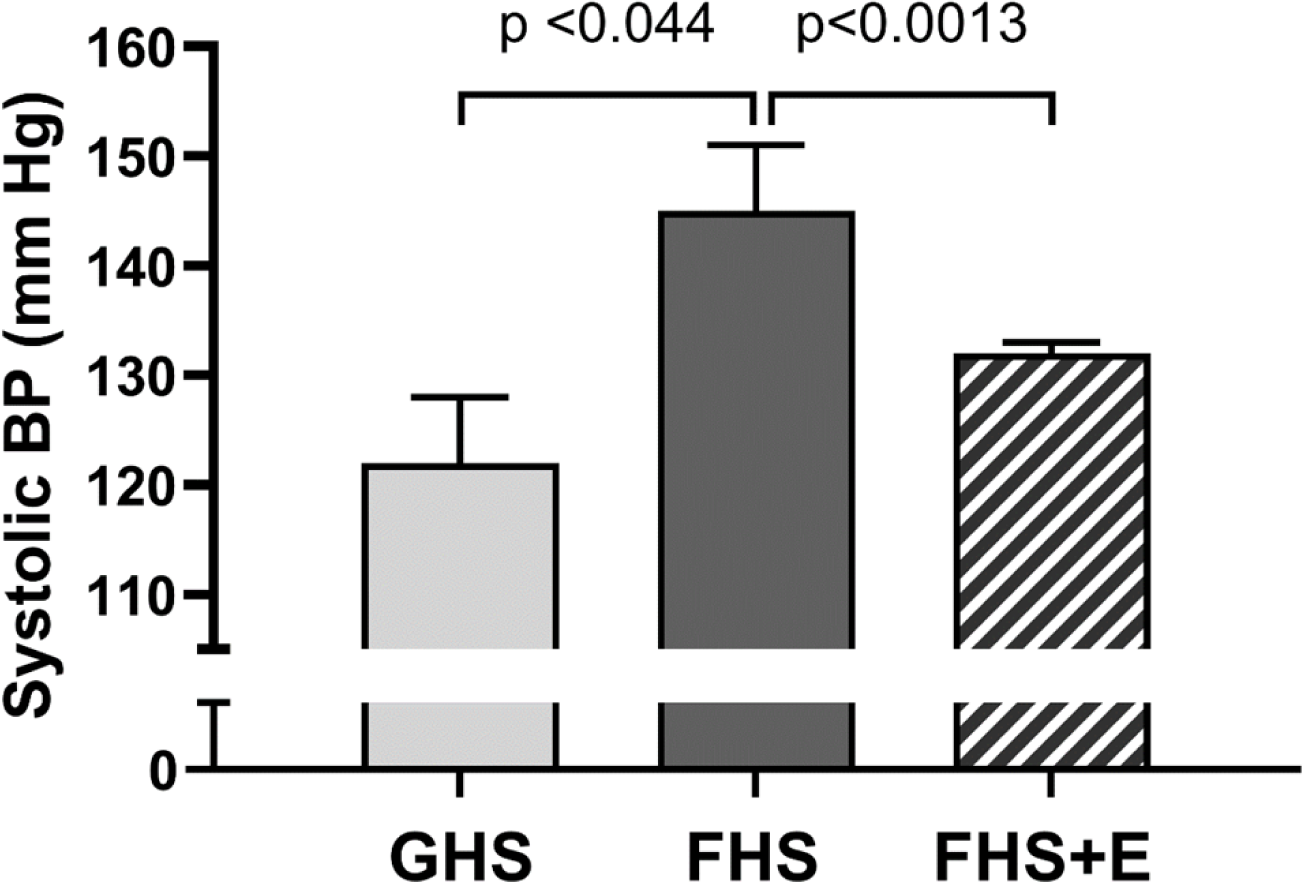
Systolic blood pressure (BP) after 7 days of dietary intervention. One-way ANOVA indicated that the means between groups were significantly different (p ≤ 0.017). Pairwise comparisons show that FHS significantly increased SBP as compared to GHS. Chronic eplerenone blunted the increase in SBP caused by FHS. There were no differences between GHS and FHS+E. Corrections for multiple comparisons were made using the Benjamini and Hochberg method, and adjusted p-values (p-Adj) are reported. Glucose high-salt (GHS; n = 6), Fructose high-salt (FHS; n=6), and FHS with coadministered eplerenone (FHS+E; n = 6).

Given that ENaC is a key channel regulated by the MR in the rat ASDN, we inquired whether blocking ENaC with the diuretic amiloride would blunt fructose-induced salt-sensitive hypertension. Therefore, we measured the effects of amiloride treatment (24 hrs) on systolic BP. Acute administration of amiloride decreased systolic BP in FHS-fed rats from 148±6 to 134±5 mm Hg, a reduction of 14±4 mm Hg (**Figure 5A**; p ≤ 0.019). In contrast, amiloride did not significantly change systolic BP in GHS (**Figure 5B**; GHS: 124±5; GHS+A: 133±6; ns.).

**Figure 5:**
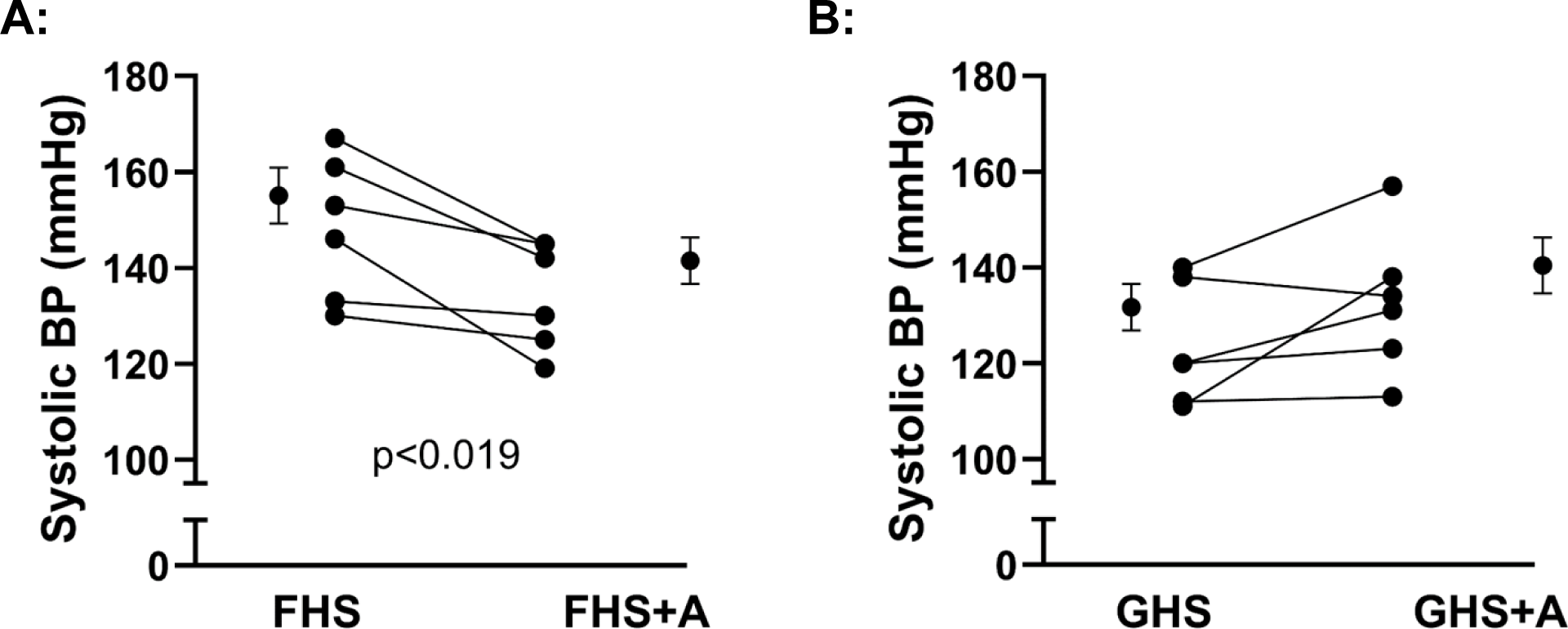
Effect of 24-hr amiloride treatment on systolic blood pressure (BP). **A:** Amiloride reduced systolic BP in rats fed fructose high-salt (FHS; p ≤ 0.019; n = 6). **B:** Amiloride did not significantly affected blood pressure in rats fed glucose high-salt (GHS; ns.; n = 6). Means, standard errors of the means, and individual data points are shown. Data were analyzed by paired t-test.

ENaC activity is partially regulated by the expression and cleavage of its subunits (39, 51, 52). Transcriptomics analysis of ENaC subunits shows that *Scnn1a*, coding for αENaC, is significantly augmented by FHS as compared to the GHS diet (Table 2). Such difference is not observed in the expression of βENaC or γENaC transcripts. However, as transcripts, expression may not correlate with protein levels, and to assess cleavage status, we also conducted Western blot analysis. We found that total αENaC expression increased by 89±14 % (1.21±0.20 normalized OD in FHS vs. 0.64±0.10 normalized OD in GHS, p ≤ 0.03), and cleaved αENaC expression increased by 47±16 % (0.85±0.08 normalized OD in FHS vs. 0.58±0.05 normalized OD in GHS, p ≤ 0.01) (**Figure 6**). Total βENaC (**Figure 7**) and total γENaC expressions were not significantly different between GHS and FHS nor was the cleaved γENaC (**Figure 8**).

**Figure 6:**
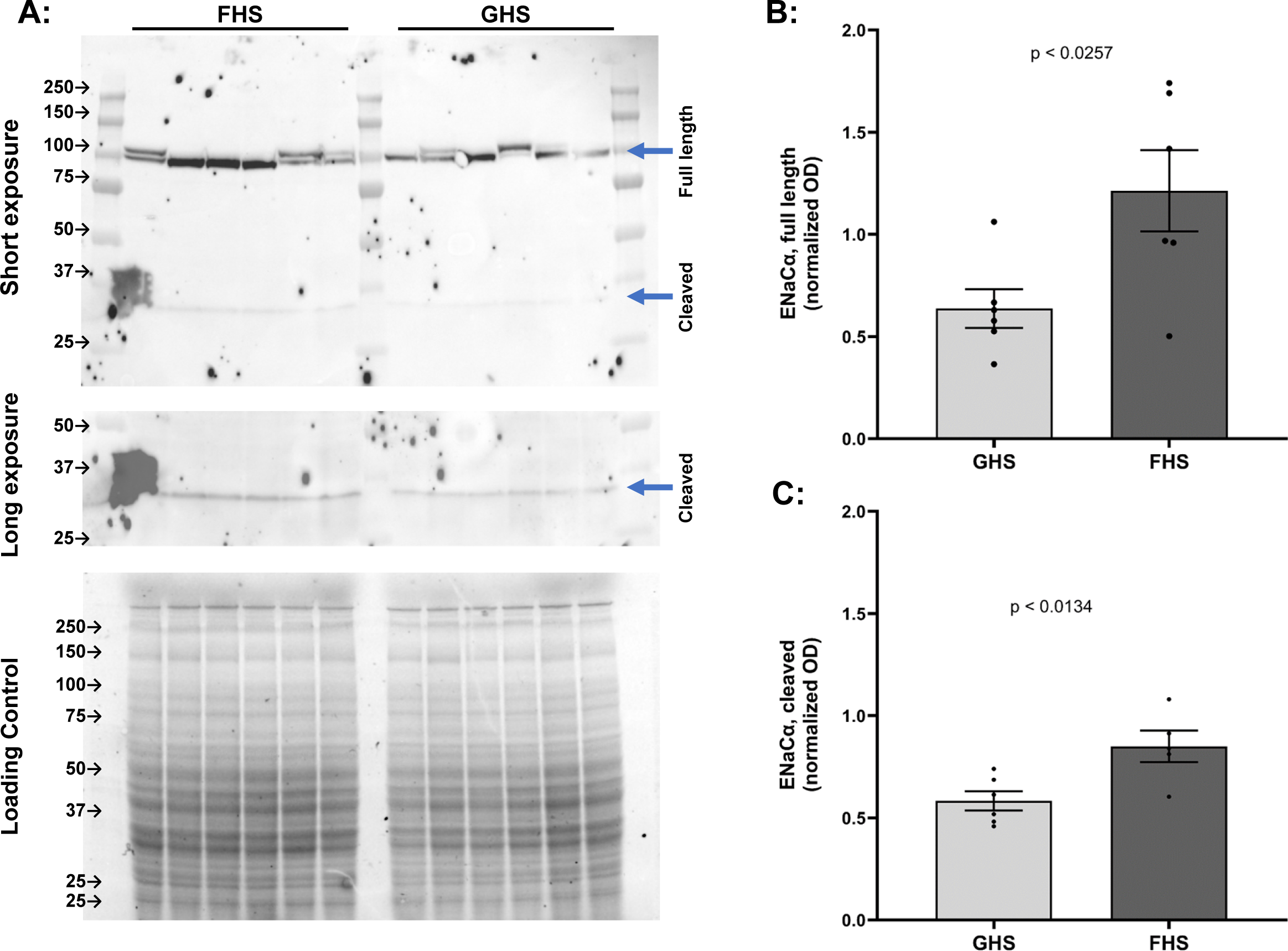
Western blot of αENaC expression in kidney cortex from Sprague Dawley rats fed fructose high-salt (FHS; n = 6) and glucose high-salt (GHS; n = 6). **A)** The upper blot shows short exposure to blot for the full length αENaC. The middle blot shows a long exposure to reveal cleaved αENaC. The bottom picture shows the total per/lane protein quantification by Stain-free technology used for normalization. **B)** Normalized optical density (OD) of full αENaC. **C)** Normalized OD of cleaved full αENaC. Unpaired t-tests were used for comparisons. Arrows indicate the lull length and cleaved αENaC as labeled.

**Figure 7:**
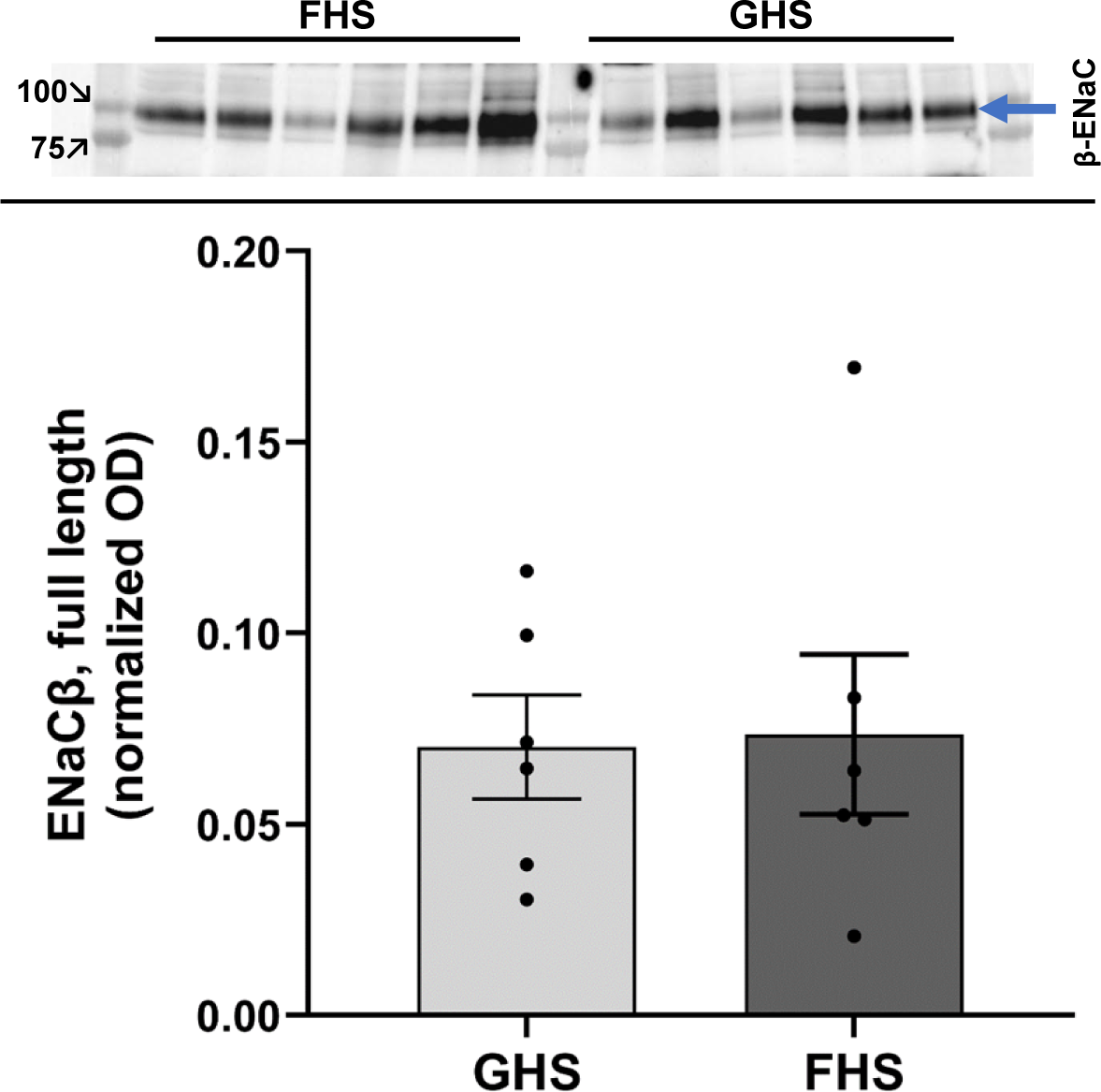
Western blot of βENaC expression in kidney cortex from rats fed fructose high-salt (FHS; n = 6) and glucose high-salt (GHS; n = 6). The blot shows βENaC expression (blue arrow). Data were compared by unpaired t-test.

**Figure 8:**
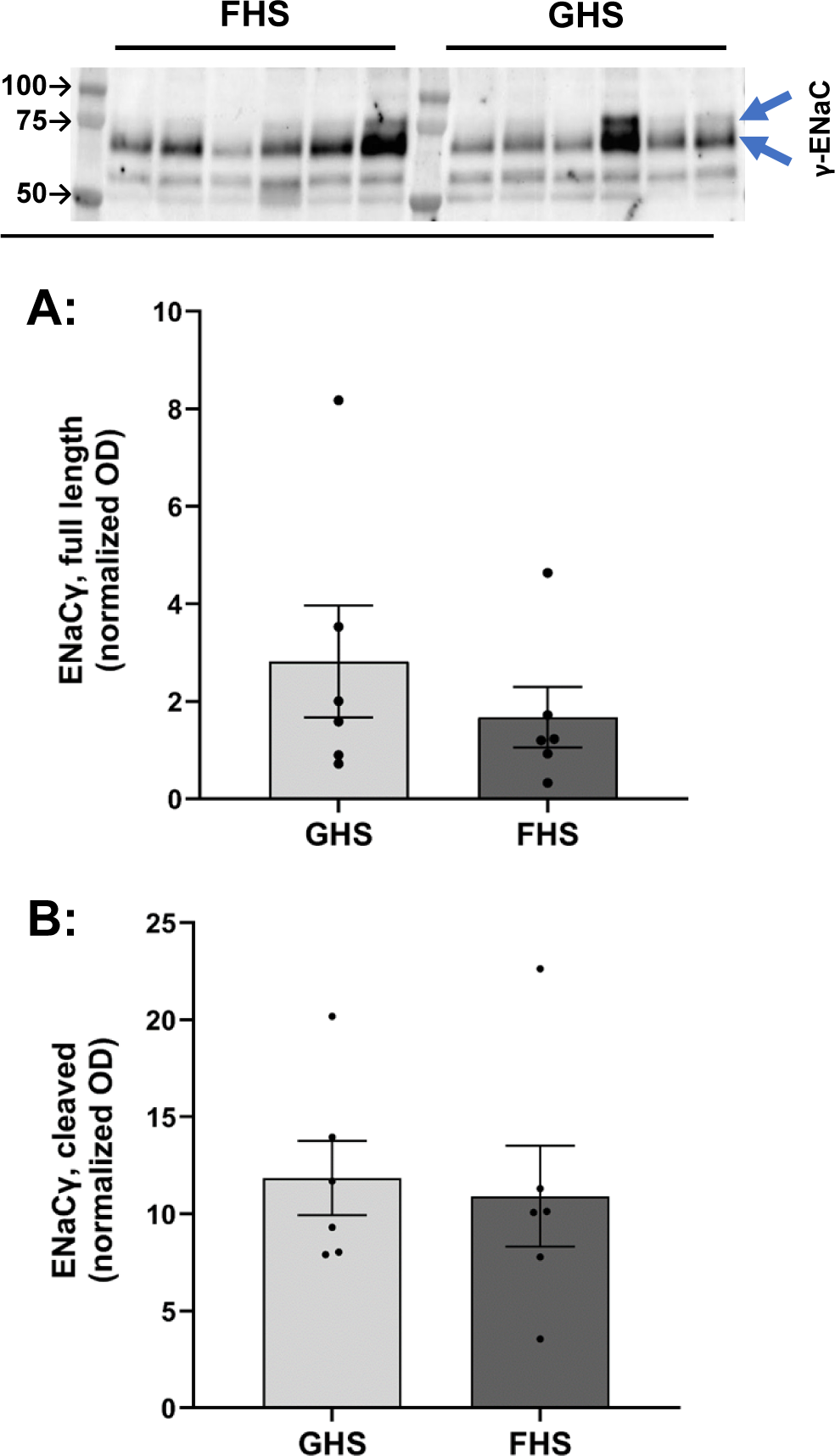
Western blot of γENaC expression in kidney cortex from rats fed fructose high-salt (FHS; n = 6) and glucose high-salt (GHS; n = 6). The blot shows full-length (upper blue arrow) and cleaved (lower blue arrow) γENaC. Densitometry is presented in **(A)** and **(B)**, respectively. Data were compared by unpaired t-test.

**Table 2:** ENaC subunits expression in relation to FHS.

| Protein | Gene Symbol | Pearson correlation with FHS |  | Wald test for differential gene expression (DESeq2) |  |  |  |
| --- | --- | --- | --- | --- | --- | --- | --- |
| | | Coefficient (r) | p-value | $\log_2\text{FC}^*$ | $\log_2\text{FC}.\text{SE}^*$ | Statistic | p-value |
| $\alpha\text{ENaC}$ | *Scnn1a* | 0.78 | 0.002 | 0.38 | 0.13 | 2.84 | 0.004 |
| $\beta\text{ENaC}$ | *Scnn1b* | 0.27 | 0.39 | 0.05 | 0.10 | 0.54 | 0.59 |
| $\gamma\text{ENaC}$ | *Scnn1g* | 0.38 | 0.22 | 0.09 | 0.10 | 0.87 | 0.38 |

\* $\log_2\text{FC}$ : log2 fold change. <sup>SE</sup>SE: standard error

## Discussion

We hypothesized that fructose-induced salt-sensitive hypertension depends in part on the abnormal activation of the mineralocorticoid receptor in the ASDN with consequent increases in ENaC expression. We found that changes in renal cortical transcriptomes induced by fructose are consistent with increased MR signaling and elevated Na^+^ reabsorption by the ASDN. Eplerenone, a MR blocker, prevented FHS from increasing BP. Plasma aldosterone concentrations in rats fed FHS were similar to those fed GHS, even though the former group was hypertensive. FHS animals also presented increased expression of 11β-HSD1, suggesting cross-activation of the MR by corticosterone. Amiloride, an ENaC inhibitor, reversed fructose-induced salt-sensitive hypertension. Furthermore, FHS augmented total and cleaved αENaC expression, while the expression of βENaC and γENaC subunits remained unchanged. In a previous study, we identified a group of coexpressed genes whose expression levels were affected by fructose. Specifically, WGCNA revealed a gene coexpression module correlated with FHS, named paleturquoise, which presented enrichment terms associated with Na^+^ reabsorption in the distal nephron. We confirmed these findings by showing that genes in the paleturquoise module are more highly expressed in the ASDN, than in proximal tubules, cortical Loop of Henle, or the glomerular epithelium. In addition, the paleturquoise genes presented a significant overlap with a transcriptional signature elicited by aldosterone in the ASDN, and blocking the MR with eplerenone prevents fructose-induced salt-sensitive elevation in BP.

The physiological activators of the MR in rodents are aldosterone and corticosterone. Aldosterone-sensitive tissues, including the ASDN, express 11β-HSD2 to prevent activation of the MR by corticosterone (53). Opposing the effects of 11β-HSD2, the actions of 11β-HSD1 could increase the local concentration of corticosterone, allowing local regulation of the MR by glucocorticoids (54). Our analysis showed that 11β-HSD1 transcripts were increased in the kidney cortex of FHS animals, suggesting locally elevated corticosterone levels. On the contrary, we found no differences in the expression of 11β-HSD2 transcripts in bulk transcriptomes. Previous studies have shown that the expression of 11β-HSD2 increases from the DCT to the CCD (55). This allows the glucocorticoid-mediated regulation of MR in the late DCT and early CNT, where ENaC activity is largely independent of aldosterone (56, 57). The opposite is true for the late CNT and cortical collecting duct (55-57).

We expected lower aldosterone levels in the hypertensive FHS group due to the physiological regulation of renin release by BP (17-19). However, the excess dietary salt bunted aldosterone in FHS and GHS alike. The systemic regulation of the renin-angiotensin-aldosterone system in FHS remains controversial. Other investigators reported decreases (58), increases (59), and lack of change (60) in plasma renin activity in animals fed fructose and high salt. Our current findings indicates that excess salt-likely override any effect of fructose in circulating aldosterone levels. Together, these data support the idea that MR genomic signaling in FHS is primarily mediated by glucocorticoids. An important limitation of these studies is that we worked with bulk transcriptomes. Thus, our current data does not resolve from which cell-type the MR transcriptional signature is coming from. Future studies using single-cell or spatially-resolved single-cell transcriptomes can clarify this issue.

Given the dependence of fructose-induced salt-sensitive hypertension on MR activation, we investigated the role of ENaC, a key channel regulated by MR in the ASDN. We found that amiloride nearly completely reversed the increase in BP caused by FHS. These are the first studies to implicate ENaC in fructose-induced salt-sensitive hypertension. ENaC activity is subjected to complex regulation, including subunit expression (61), membrane insertion and internalization, and cleavage of the α- and γ-subunits with release of embedded inhibitory tracts. Proteolytic cleavage of the α-and γ-subunits occurs during intracellular processing by the protease furin, and at the luminal membrane by a variety of cell surface or urinary proteases (52, 62, 63). To understand if FHS activates ENaC we investigated ENaC transcripts in bulk RNAseq transcriptomes, and showed that transcripts for αENaC were increased by FHS, but not those for γENaC or βENaC. These data are consistent with other reports showing that αENaC is the primary effector of MR activation in rodents (61, 64). Then, we assessed ENaC protein expression and cleavage by Western blots. Similar to the transcriptomic data, we found that the total expression of αENaC was increased by FHS, while γENaC and βENaC levels did not change. Interestingly, even though cleaved αENaC was slightly increased, the cleavage ratios of either αENaC or γENaC were not changed by FHS. This, lack of an increase in cleaved ENaC subunits suggests that additional activation by urinary proteases does not play a major role in fructose-induced salt-sensitive hypertension. However, it should be noted that an increase in total expression of cell surface ENaC, in association with the increase in αENaC expression, likely contributes to an increase in ENaC activity and in BP. Finally, an activation of ENaC due to the increase in luminal flow caused by the high-salt diet is unlikely to explain differences in BP in FHS, as it will be equally enhanced by GHS.

We previously showed that the bulk of filtered fructose is reabsorbed and metabolized in proximal tubules (44, 65), where it increased Na^+^ reabsorption in response to Ang II (29, 30, 66). In addition, others have reported fructose can also enhance Na^+^ reabsorption in thick ascending limbs (67) and the DCT (41). The present work extends these findings to the ASDN, further highlighting that fructose causes dysregulation of homeostatic mechanisms in multiple parts of the nephron.

The data presented here show that, unlike glucose, fructose causes salt-sensitive increases in BP within a week. This is consistent with our previous reports administering 20% fructose as solid diet (22, 31) or in the drinking water (29, 49), as well as publications from other investigators using similar diet regimes (59, 60, 68). Importantly, the amount of fructose used in these studies is consumed by more than 17 million Americans, and likely contributes to salt-sensitive hypertension in the US population. Because we were interested in the causes of the elevated BP, we only studied rats early in the development of hypertension, *i.e.* at 7-10 days. In this time frame, there is no overt renal damage, and rats are euglycemic (29). Our model differs from others used to induce metabolic syndrome, like those using greater amounts of fructose chronically (69-71), or protocols in which rats were first fed a fructose low-salt diet and then switched to FHS (59, 60, 66, 68, 72). It is critical to note, that while the data presented here support the hypothesis that fructose-induced salt-sensitive hypertension depends in part on MR activation by the ASDN, the mechanisms involved are less clear and have limitations. Multiple lines of evidence pointed us to ENaC, but transcript or protein levels are not necessarily indicative of ENaC activity, and we did not perform membrane trafficking studies or measurements of this channel by patch clamping. Whole-cell patch clamping studies between the DCT2-early CNT as compared to late CNT-CCD, would also help clarify whether the increase in ENaC depends in aldosterone or glucocorticoids (55, 57).

In summary, our study provides new insights into the pathophysiology of fructose-induced salt-sensitive hypertension, suggesting a pivotal role of MR and ENaC.

## Data Availability Statement

The raw sequencing files utilized in this study are openly available from xxxTBD and xxxTBD as described in the Methods section for each dataset. The code used for all bioinformatics experiments and to generate the corresponding figures is available at https://github.com/AgustinGonvi/TBD.

## Author Contributions

R.Z., S.S., J.L.G., T.K., and A.G-V. conceptualized the study; R.Z., S.S., D.A.J., N.K., A.B., R.S., C.Q., B.K., and A.G-V. performed experiments and collected data; R.Z., S.S., D.A.J., J.L.G., and A.G-V. curated and analyzed data; R.Z., S.S., J.L.G., B.R.F., T.K., and A.G-V. interpreted results; R.Z., J.L.G., and A.G-V. prepared the manuscript and figures. All authors provided critical feedback, and have read and agreed to the final version of the manuscript.

## Acknowledgements

Authors would like to thank Simone Edelheit and the Genomics Core at CWRU School of Medicine for their support in library preparation and sequencing, and to Johannes Loffing for providing the anti αENaC antibody.

---

**Supplemental Table 1:**
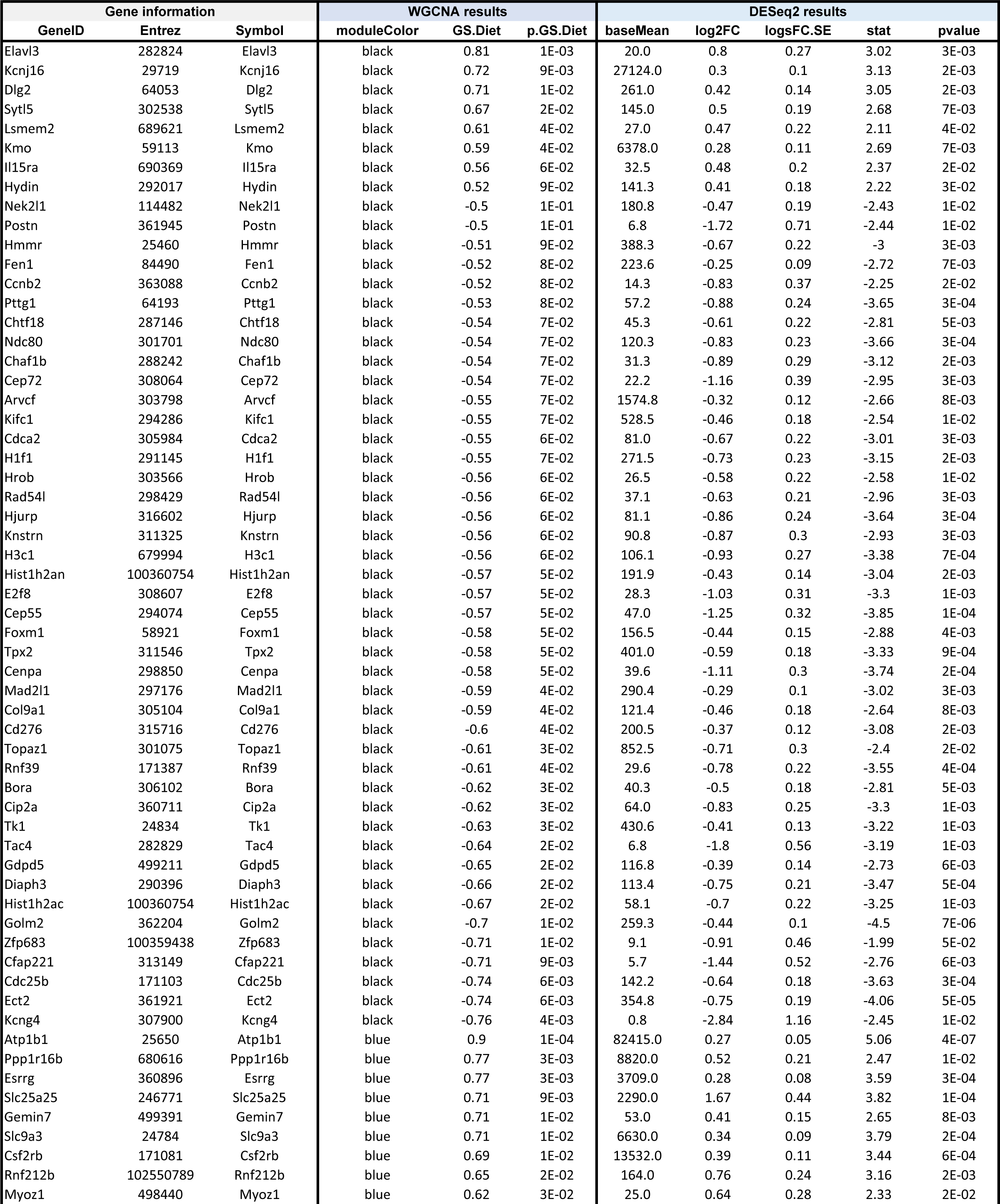
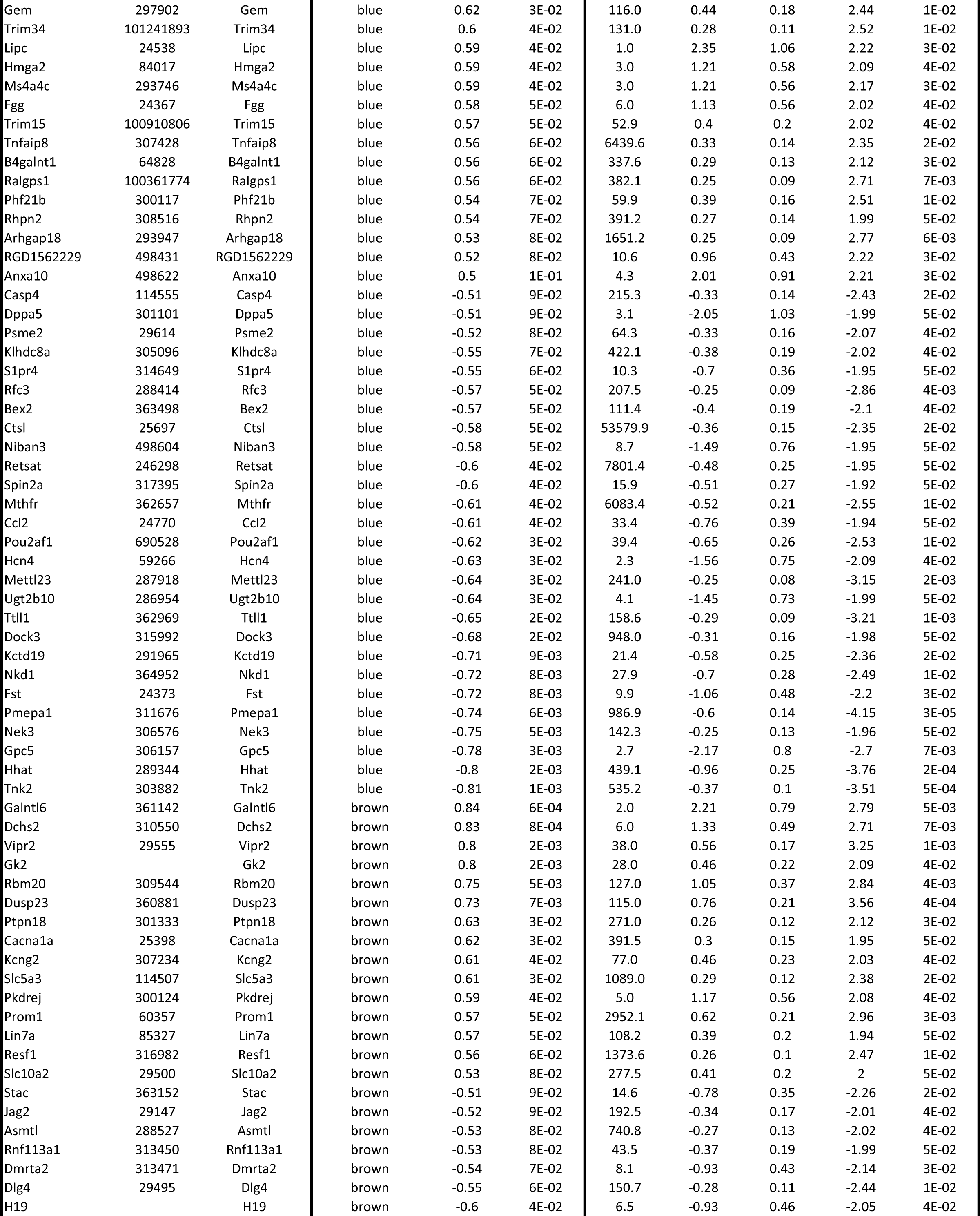
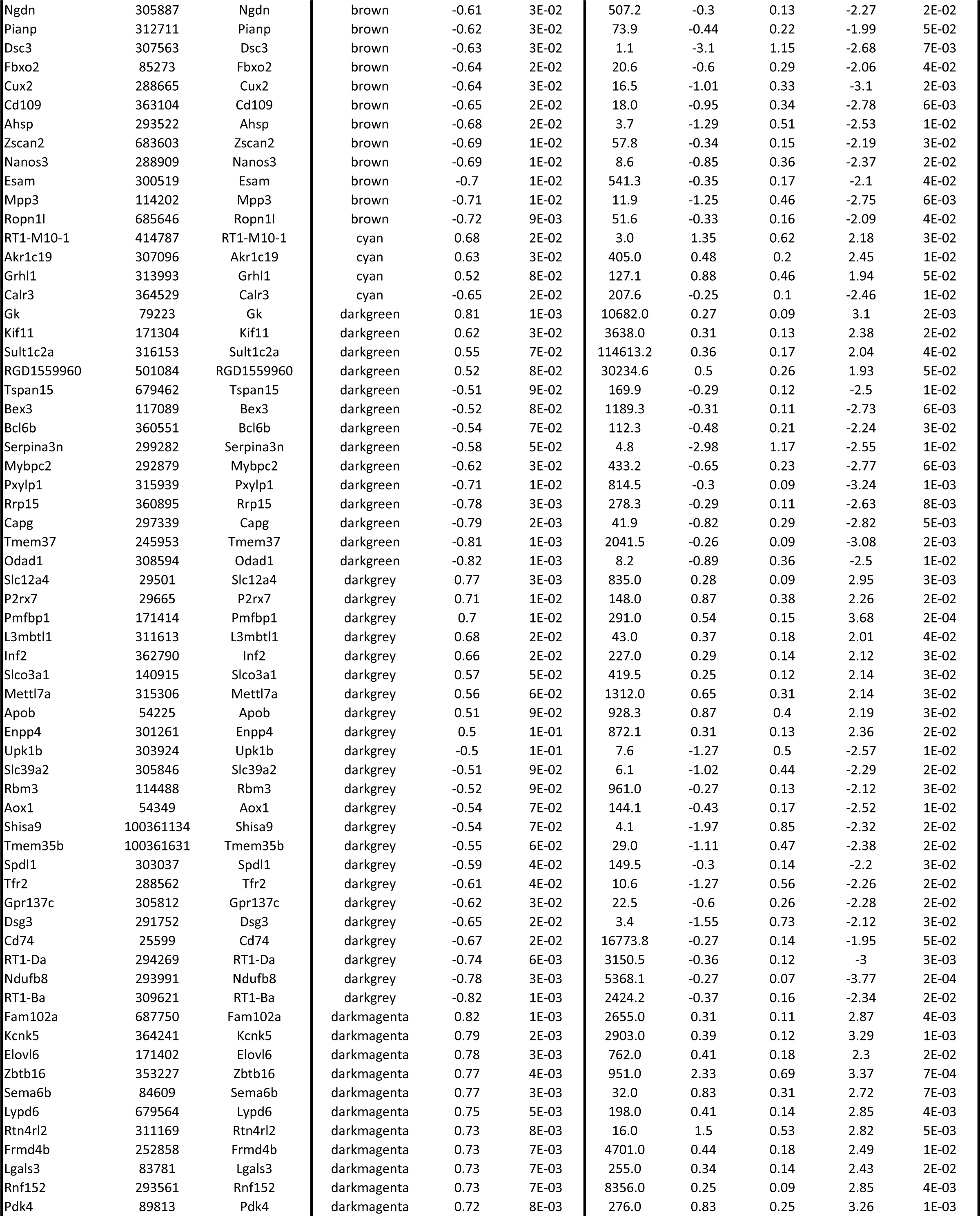
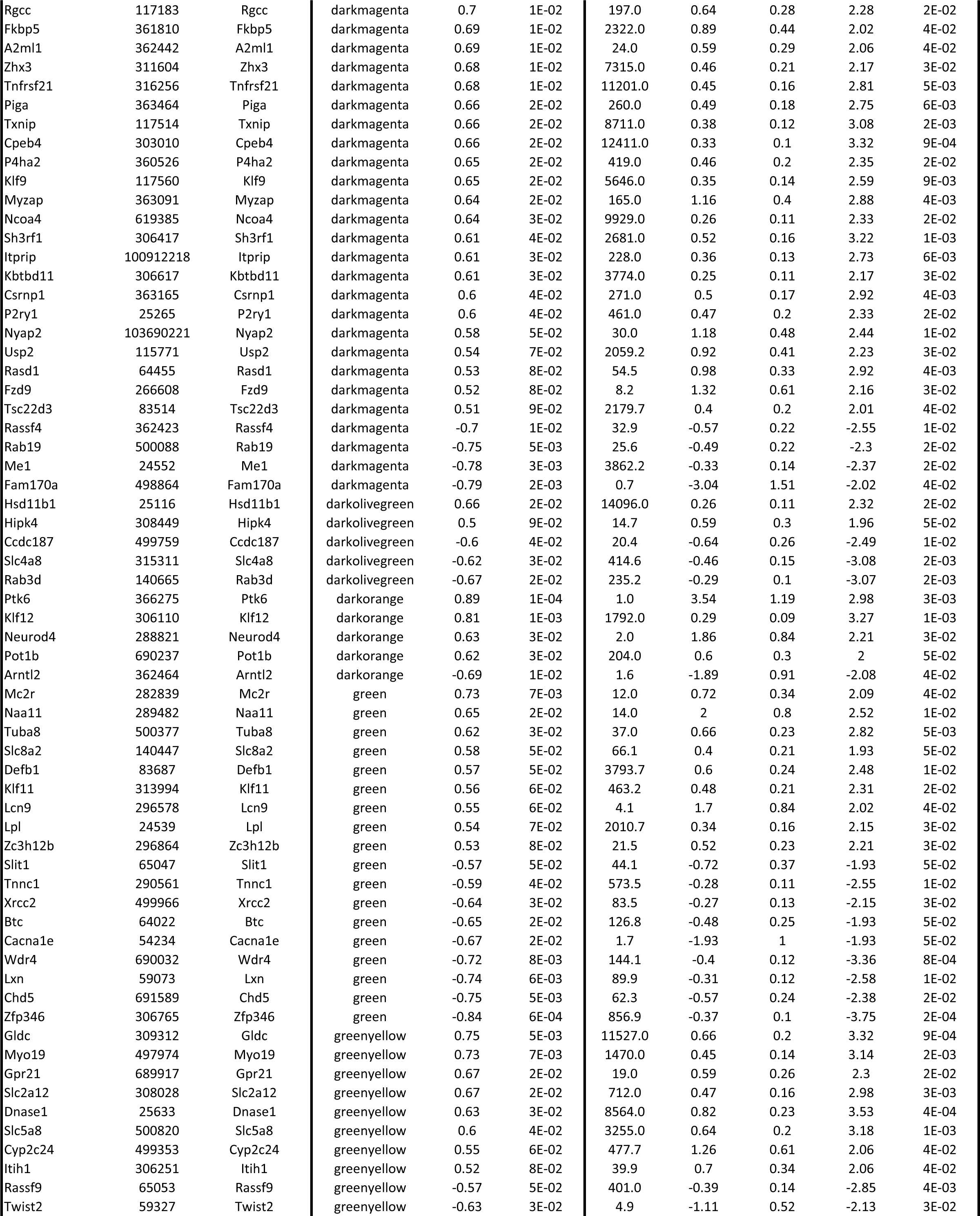
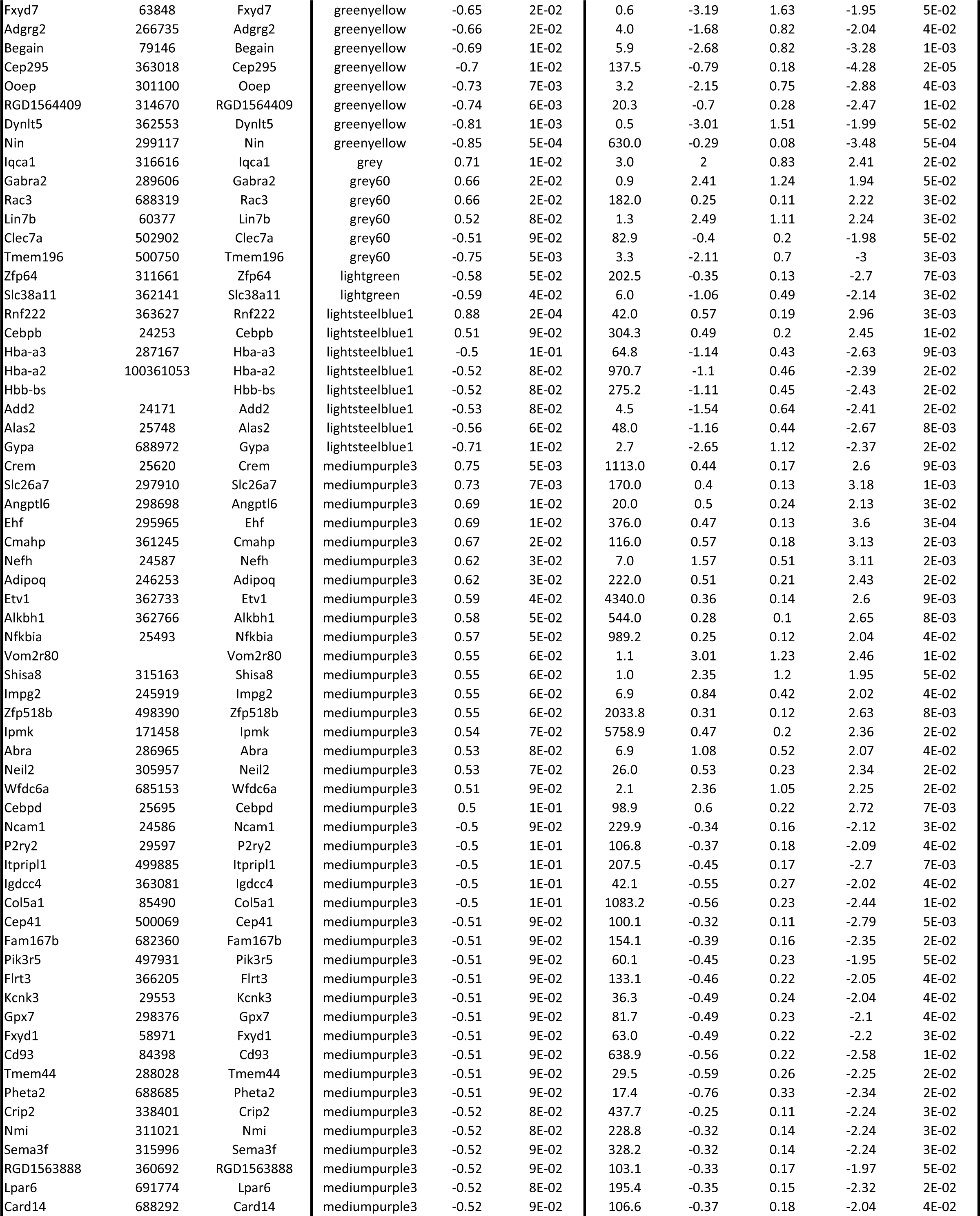
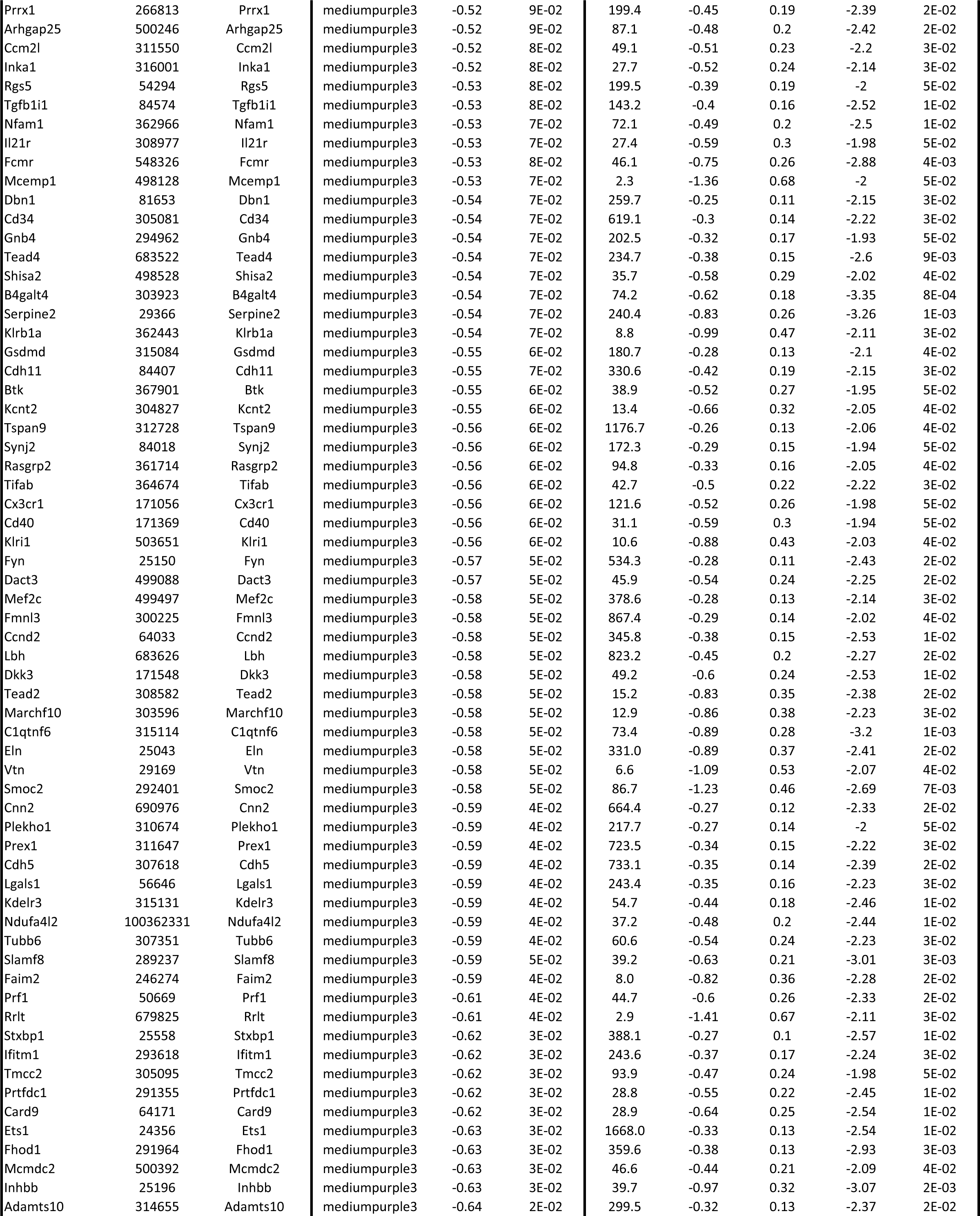
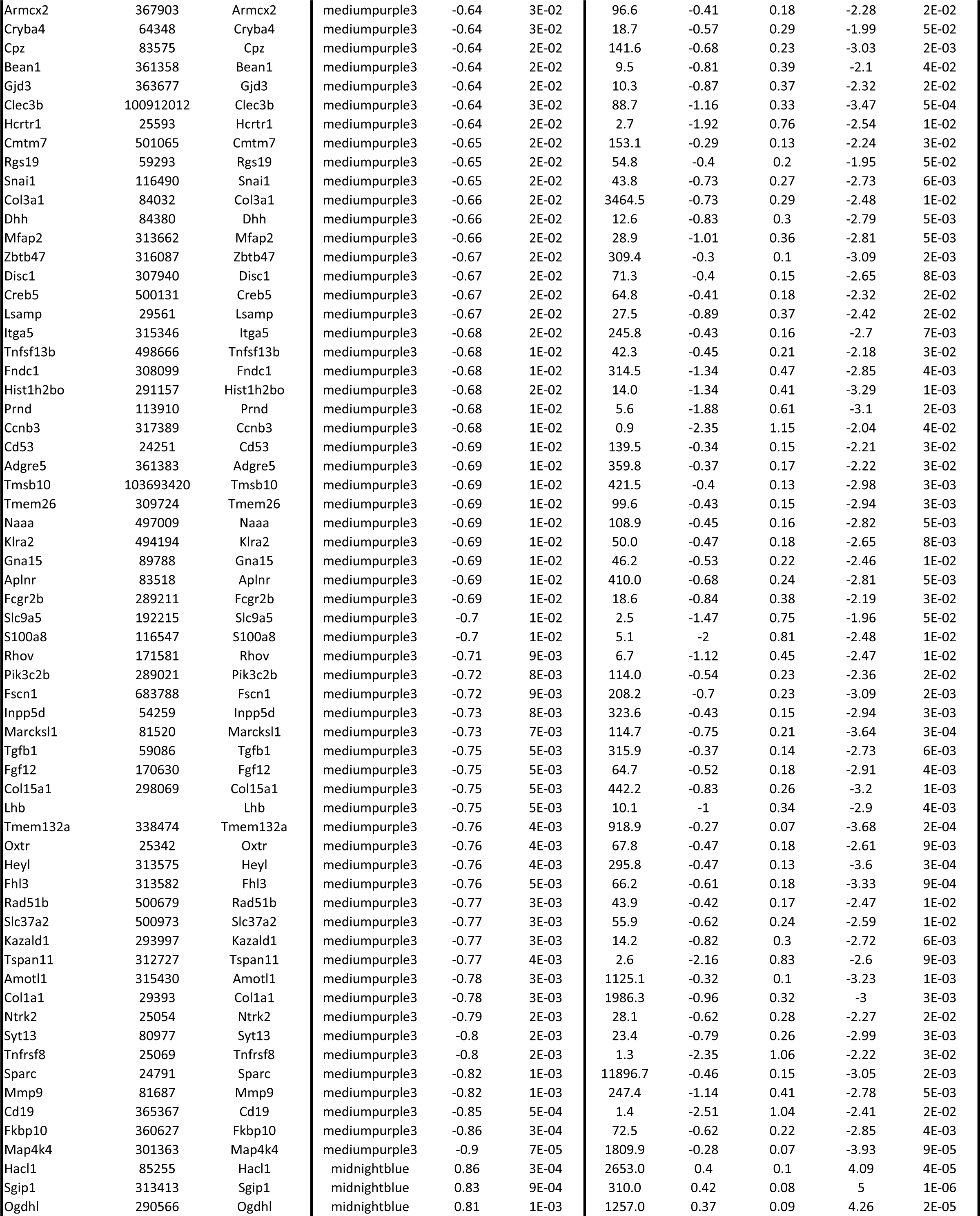
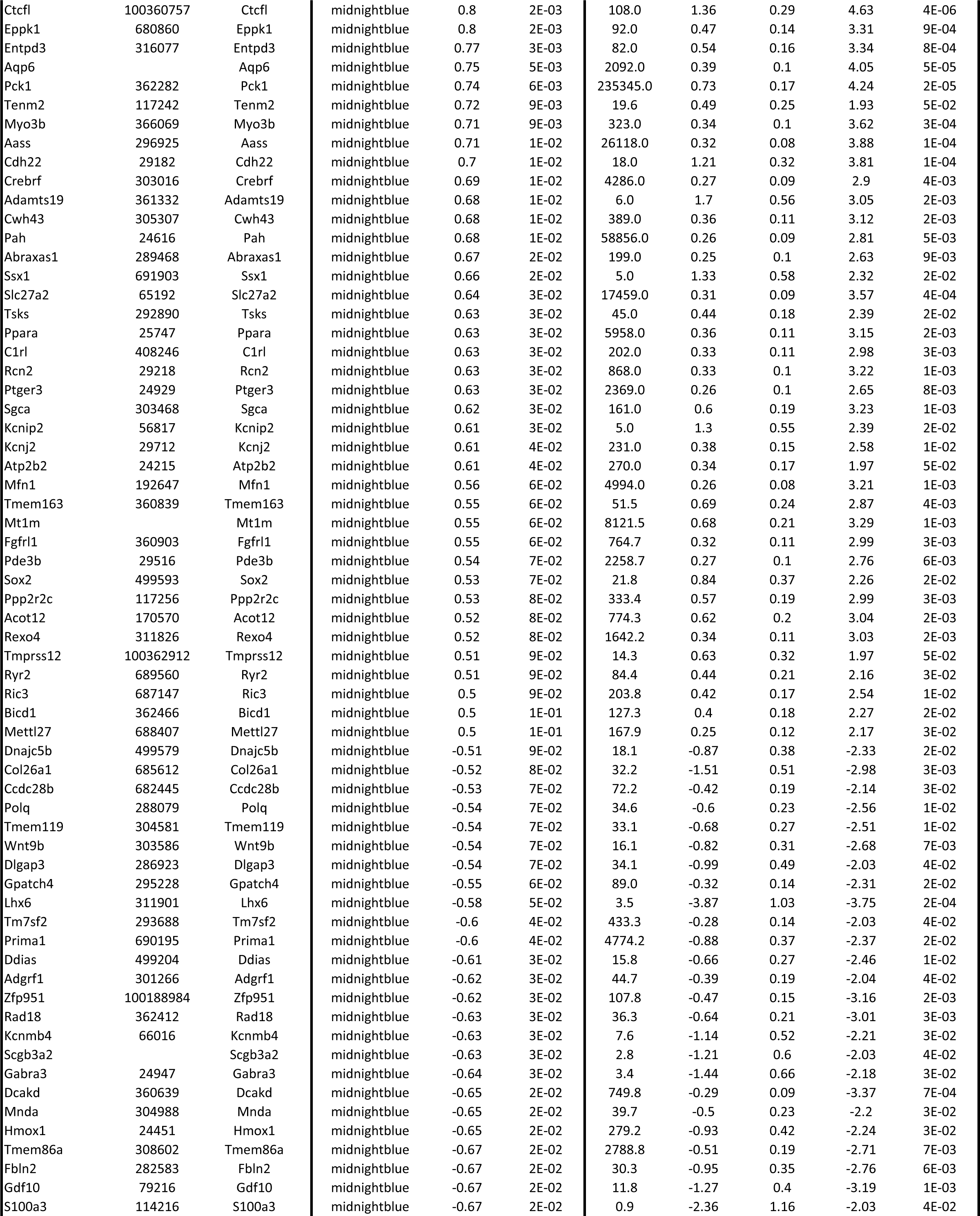
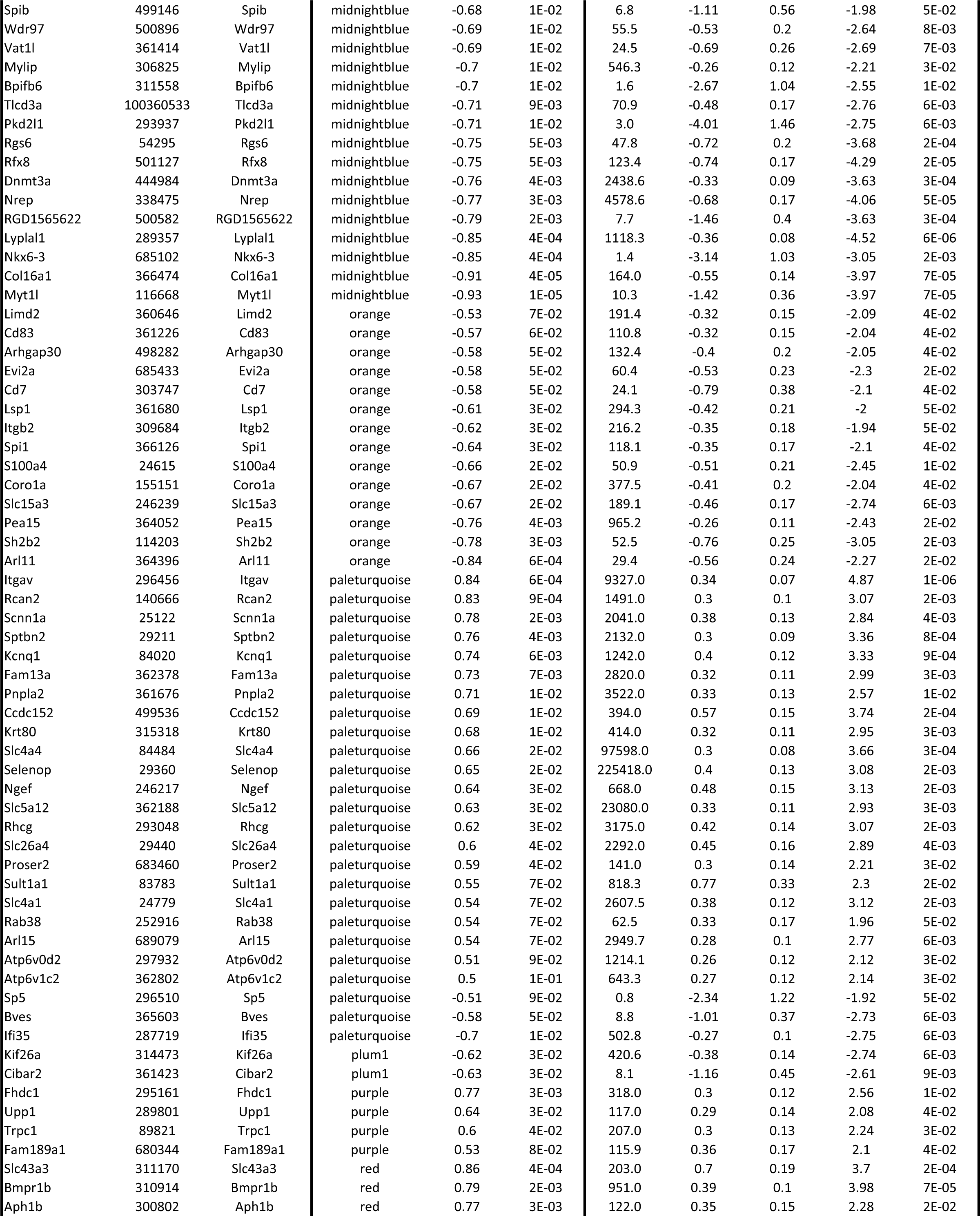
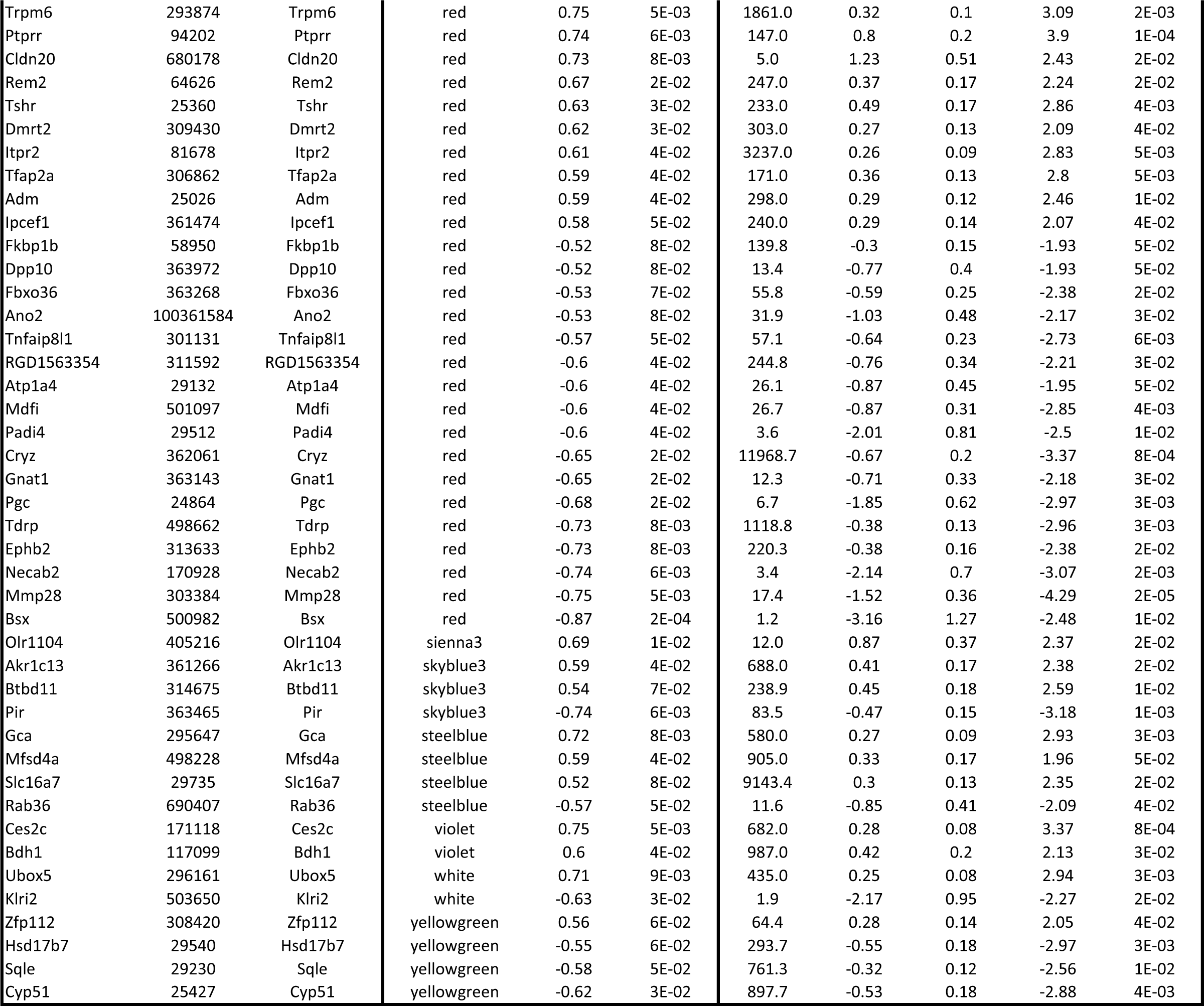

## References

1. Lim SS, Vos T, Flaxman AD, Danaei G, Shibuya K, Adair-Rohani H, Amann M, Anderson HR, Andrews KG, Aryee M, Atkinson C, Bacchus LJ, Bahalim AN, Balakrishnan K, Balmes J, Barker-Collo S, Baxter A, Bell ML, Blore JD, Blyth F, Bonner C, Borges G, Bourne R, Boussinesq M, Brauer M, Brooks P, Bruce NG, Brunekreef B, Bryan-Hancock C, Bucello C, Buchbinder R, Bull F, Burnett RT, Byers TE, Calabria B, Carapetis J, Carnahan E, Chafe Z, Charlson F, Chen H, Chen JS, Cheng AT, Child JC, Cohen A, Colson KE, Cowie BC, Darby S, Darling S, Davis A, Degenhardt L, Dentener F, Des Jarlais DC, Devries K, Dherani M, Ding EL, Dorsey ER, Driscoll T, Edmond K, Ali SE, Engell RE, Erwin PJ, Fahimi S, Falder G, Farzadfar F, Ferrari A, Finucane MM, Flaxman S, Fowkes FG, Freedman G, Freeman MK, Gakidou E, Ghosh S, Giovannucci E, Gmel G, Graham K, Grainger R, Grant B, Gunnell D, Gutierrez HR, Hall W, Hoek HW, Hogan A, Hosgood HD, 3rd, Hoy D, Hu H, Hubbell BJ, Hutchings SJ, Ibeanusi SE, Jacklyn GL, Jasrasaria R, Jonas JB, Kan H, Kanis JA, Kassebaum N, Kawakami N, Khang YH, Khatibzadeh S, Khoo JP, Kok C, Laden F, Lalloo R, Lan Q, Lathlean T, Leasher JL, Leigh J, Li Y, Lin JK, Lipshultz SE, London S, Lozano R, Lu Y, Mak J, Malekzadeh R, Mallinger L, Marcenes W, March L, Marks R, Martin R, McGale P, McGrath J, Mehta S, Mensah GA, Merriman TR, Micha R, Michaud C, Mishra V, Mohd Hanafiah K, Mokdad AA, Morawska L, Mozaffarian D, Murphy T, Naghavi M, Neal B, Nelson PK, Nolla JM, Norman R, Olives C, Omer SB, Orchard J, Osborne R, Ostro B, Page A, Pandey KD, Parry CD, Passmore E, Patra J, Pearce N, Pelizzari PM, Petzold M, Phillips MR, Pope D, Pope CA, 3rd, Powles J, Rao M, Razavi H, Rehfuess EA, Rehm JT, Ritz B, Rivara FP, Roberts T, Robinson C, Rodriguez-Portales JA, Romieu I, Room R, Rosenfeld LC, Roy A, Rushton L, Salomon JA, Sampson U, Sanchez-Riera L, Sanman E, Sapkota A, Seedat S, Shi P, Shield K, Shivakoti R, Singh GM, Sleet DA, Smith E, Smith KR, Stapelberg NJ, Steenland K, Stockl H, Stovner LJ, Straif K, Straney L, Thurston GD, Tran JH, Van Dingenen R, van Donkelaar A, Veerman JL, Vijayakumar L, Weintraub R, Weissman MM, White RA, Whiteford H, Wiersma ST, Wilkinson JD, Williams HC, Williams W, Wilson N, Woolf AD, Yip P, Zielinski JM, Lopez AD, Murray CJ, Ezzati M, AlMazroa MA, and Memish ZA. A comparative risk assessment of burden of disease and injury attributable to 67 risk factors and risk factor clusters in 21 regions, 1990-2010: a systematic analysis for the Global Burden of Disease Study 2010. Lancet 380: 2224-2260, 2012.

2. Joynt Maddox KE, Elkind MSV, Aparicio HJ, Commodore-Mensah Y, de Ferranti SD, Dowd WN, Hernandez AF, Khavjou O, Michos ED, Palaniappan L, Penko J, Poudel R, Roger VL, Kazi DS, and American Heart A. Forecasting the Burden of Cardiovascular Disease and Stroke in the United States Through 2050-Prevalence of Risk Factors and Disease: A Presidential Advisory From the American Heart Association. Circulation 150: e65–e88, 2024.

3. Weinberger MH. Salt sensitivity of blood pressure in humans. Hypertension 27: 481–490, 1996.

4. Sullivan JM. Salt sensitivity. Definition, conception, methodology, and long-term issues. Hypertension 17: I61–68, 1991.

5. Peters RM, and Flack JM. Salt sensitivity and hypertension in African Americans: implications for cardiovascular nurses. Prog Cardiovasc Nurs 15: 138–144, 2000.

6. Svetkey LP, McKeown SP, and Wilson AF. Heritability of salt sensitivity in black Americans. Hypertension 28: 854–858, 1996.

7. Luft FC, Miller JZ, Grim CE, Fineberg NS, Christian JC, Daugherty SA, and Weinberger MH. Salt Sensitivity and Resistance of Blood-Pressure - Age and Race as Factors in Physiological-Responses. Hypertension 17: I102–I108, 1991.

8. Guyton AC. Physiologic regulation of arterial pressure. Am J Cardiol 8: 401–407, 1961.

9. Roman RJ. Abnormal renal hemodynamics and pressure-natriuresis relationship in Dahl salt-sensitive rats. Am J Physiol 251: F57–65, 1986.

10. Marriott BP, Olsho L, Hadden L, and Connor P. Intake of added sugars and selected nutrients in the United States, National Health and Nutrition Examination Survey (NHANES) 2003-2006. Critical reviews in food science and nutrition 50: 228-258, 2010.

11. Vos MB, Kimmons JE, Gillespie C, Welsh J, and Blanck HM. Dietary fructose consumption among US children and adults: the Third National Health and Nutrition Examination Survey. Medscape journal of medicine 10: 160, 2008.

12. Ogden CL, Kit BK, Carroll MD, and Park S. Consumption of sugar drinks in the United States, 2005-2008. NCHS data brief 1-8, 2011.

13. Hwang IS, Ho H, Hoffman BB, and Reaven GM. Fructose-induced insulin resistance and hypertension in rats. Hypertension 10: 512–516, 1987.

14. Nguyen S, Choi HK, Lustig RH, and Hsu CY. Sugar-sweetened beverages, serum uric acid, and blood pressure in adolescents. The Journal of pediatrics 154: 807–813, 2009.

15. Jalal DI, Smits G, Johnson RJ, and Chonchol M. Increased fructose associates with elevated blood pressure. J Am Soc Nephrol 21: 1543–1549, 2010.

16. Sechi LA. Mechanisms of insulin resistance in rat models of hypertension and their relationships with salt sensitivity. Journal of hypertension 17: 1229–1237, 1999.

17. Nishimoto Y, Tomida T, Matsui H, Ito T, and Okumura K. Decrease in renal medullary endothelial nitric oxide synthase of fructose-fed, salt-sensitive hypertensive rats. Hypertension 40: 190–194, 2002.

18. Brown IJ, Stamler J, Van Horn L, Robertson CE, Chan Q, Dyer AR, Huang CC, Rodriguez BL, Zhao L, Daviglus ML, Ueshima H, Elliott P, International Study of MM, and Blood Pressure Research G. Sugar-sweetened beverage, sugar intake of individuals, and their blood pressure: international study of macro/micronutrients and blood pressure. Hypertension 57: 695–701, 2011.

19. Grasser EK, Dulloo A, and Montani JP. Cardiovascular responses to the ingestion of sugary drinks using a randomised cross-over study design: Does glucose attenuate the blood pressure-elevating effect of fructose? Br J Nutr 112: 183–192, 2014.

20. Dhingra R, Sullivan L, Jacques PF, Wang TJ, Fox CS, Meigs JB, D’Agostino RB, Gaziano JM, and Vasan RS. Soft drink consumption and risk of developing cardiometabolic risk factors and the metabolic syndrome in middle-aged adults in the community. Circulation 116: 480–488, 2007.

21. Dai S, and McNeill JH. Fructose-induced hypertension in rats is concentration- and duration-dependent. Journal of pharmacological and toxicological methods 33: 101–107, 1995.

22. Forester BR, Brostek A, Schuhler B, Gonzalez-Vicente A, and Garvin JL. Angiotensin II-stimulated proximal nephron superoxide production and fructose-induced salt-sensitive hypertension. American journal of physiology Renal physiology 326: F249–F256, 2024.

23. Ferrario CM, Groban L, Wang H, Sun X, VonCannon JL, Wright KN, and Ahmad S. The renin-angiotensin system biomolecular cascade: a 2022 update of newer insights and concepts. Kidney Int Suppl (2011) 12: 36–47, 2022.

24. Mirabito Colafella KM, and Danser AHJ. Recent Advances in Angiotensin Research. Hypertension 69: 994–999, 2017.

25. Nguyen Dinh Cat A, and Touyz RM. A new look at the renin-angiotensin system--focusing on the vascular system. Peptides 32: 2141-2150, 2011.

26. Garvin JL, and Beierwaltes WH. Response of proximal tubules to angiotensin II changes during maturation. Hypertension 31: 415–420, 1998.

27. Harris PJ. Regulation of proximal tubule function by angiotensin. Clin Exp Pharmacol Physiol 19: 213–222, 1992.

28. Yingst DR, Massey KJ, Rossi NF, Mohanty MJ, and Mattingly RR. Angiotensin II directly stimulates activity and alters the phosphorylation of Na-K-ATPase in rat proximal tubule with a rapid time course. American journal of physiology Renal physiology 287: F713–721, 2004.

29. Gonzalez-Vicente A, Hong NJ, Yang N, Cabral PD, Berthiaume JM, Dominici FP, and Garvin JL. Dietary Fructose Increases the Sensitivity of Proximal Tubules to Angiotensin II in Rats Fed High-Salt Diets. Nutrients 10: 2018.

30. Gonzalez-Vicente A, Cabral PD, Hong NJ, Asirwatham J, Yang N, Berthiaume JM, Dominici FP, and Garvin JL. Dietary Fructose Enhances the Ability of Low Concentrations of Angiotensin II to Stimulate Proximal Tubule Na(+) Reabsorption. Nutrients 9: 2017.

31. Forester BR, Zhang R, Schuhler B, Brostek A, Gonzalez-Vicente A, and Garvin JL. Knocking Out Sodium Glucose-Linked Transporter 5 Prevents Fructose-Induced Renal Oxidative Stress and Salt-Sensitive Hypertension. Hypertension 81: 1296–1307, 2024.

32. Yang N, Gonzalez-Vicente A, and Garvin JL. Angiotensin II-induced superoxide and decreased glutathione in proximal tubules: effect of dietary fructose. American journal of physiology Renal physiology 318: F183–F192, 2020.

33. Fredlund P, Saltman S, and Catt KJ. Aldosterone production by isolated adrenal glomerulosa cells: stimulation by physiological concentrations of angiotensin II. Endocrinology 97: 1577–1586, 1975.

34. Rossier BC, Baker ME, and Studer RA. Epithelial sodium transport and its control by aldosterone: the story of our internal environment revisited. Physiological reviews 95: 297–340, 2015.

35. Wynne BM, Mistry AC, Al-Khalili O, Mallick R, Theilig F, Eaton DC, and Hoover RS. Aldosterone Modulates the Association between NCC and ENaC. Scientific reports 7: 4149, 2017.

36. Kristensen M, Fenton RA, and Poulsen SB. Dissecting the Effects of Aldosterone and Hypokalemia on the Epithelial Na(+) Channel and the NaCl Cotransporter. Front Physiol 13: 800055, 2022.

37. Poulsen SB, Limbutara K, Fenton RA, Pisitkun T, and Christensen BM. RNA sequencing of kidney distal tubule cells reveals multiple mediators of chronic aldosterone action. Physiological genomics 50: 343–354, 2018.

38. Hager H, Kwon TH, Vinnikova AK, Masilamani S, Brooks HL, Frokiaer J, Knepper MA, and Nielsen S. Immunocytochemical and immunoelectron microscopic localization of alpha-, beta-, and gamma-ENaC in rat kidney. American journal of physiology Renal physiology 280: F1093–1106, 2001.

39. Kleyman TR, and Eaton DC. Regulating ENaC’s gate. Am J Physiol Cell Physiol 318: C150–C162, 2020.

40. Anand D, Hummler E, and Rickman OJ. ENaC activation by proteases. Acta Physiol (Oxf) 235: e13811, 2022.

41. Bahena-Lopez JP, Rojas-Vega L, Chavez-Canales M, Bazua-Valenti S, Bautista-Perez R, Lee JH, Madero M, Vazquez-Manjarrez N, Alquisiras-Burgos I, Hernandez-Cruz A, Castaneda-Bueno M, Ellison DH, and Gamba G. Glucose/Fructose Delivery to the Distal Nephron Activates the Sodium-Chloride Cotransporter via the Calcium-Sensing Receptor. J Am Soc Nephrol 34: 55–72, 2023.

42. Gonzalez-Vicente A, Hong NJ, Cabral PD, and Garvin JL. A Moderately-Enriched Fructose Diet Increases The Sensitivity Of The Proximal Nephron To Angiotensin II. In: Hypertension 2014, p. A304-A304.

43. Gonzalez-Vicente A, Garvin JL, and Hopfer U. Transcriptome signature for dietary fructose-specific changes in rat renal cortex: A quantitative approach to physiological relevance. PLoS One 13: e0201293, 2018.

44. Zhang R, Jadhav DA, Kim N, Kramer B, Gonzalez-Vicente A, and Kidney Precision Medicine P. Profiling Cell Heterogeneity and Fructose Transporter Expression in the Rat Nephron by Integrating Single-Cell and Microdissected Tubule Segment Transcriptomes. Int J Mol Sci 25: 2024.

45. Zhang B, and Horvath S. A general framework for weighted gene co-expression network analysis. Stat Appl Genet Mol Biol 4: Article17, 2005.

46. Langfelder P, and Horvath S. WGCNA: an R package for weighted correlation network analysis. BMC Bioinformatics 9: 559, 2008.

47. Lake BB, and Zhang K. Isolation of single nuclei from solid tissues Kidney Precision Medicine Project. https://www.protocols.io/view/isolation-of-single-nuclei-from-solid-tissues-ufketkw.pdf. [June 20, 2023].

48. Knepper MA. Epithelial Systems Biology Laboratory https://esbl.nhlbi.nih.gov/Databases/KSBP2/. [11/10/2023.

49. Brostek A, Hong NJ, Zhang R, Forester BR, Barmore LE, Kaydo L, Kluge N, Smith C, Garvin JL, and Gonzalez-Vicente A. Independent effects of sex and stress on fructose-induced salt-sensitive hypertension. Physiological reports 10: e15489, 2022.

50. Sorensen MV, Grossmann S, Roesinger M, Gresko N, Todkar AP, Barmettler G, Ziegler U, Odermatt A, Loffing-Cueni D, and Loffing J. Rapid dephosphorylation of the renal sodium chloride cotransporter in response to oral potassium intake in mice. Kidney Int 83: 811–824, 2013.

51. Mutchler SM, Kirabo A, and Kleyman TR. Epithelial Sodium Channel and Salt-Sensitive Hypertension. Hypertension 77: 759–767, 2021.

52. Kashlan OB, Wang XP, Sheng S, and Kleyman TR. Epithelial Na (+) Channels Function as Extracellular Sensors. Compr Physiol 14: 1–41, 2024.

53. Ueda K, and Shimosawa T. Regulating distal tubule functions and salt sensitivity. American journal of physiology Renal physiology 2024.

54. Jia G, Lastra G, Bostick BP, LahamKaram N, Laakkonen JP, Yla-Herttuala S, and Whaley-Connell A. The Mineralocorticoid Receptors in diabetic kidney disease. American journal of physiology Renal physiology 2024.

55. Nesterov V, Bertog M, Canonica J, Hummler E, Coleman R, Welling PA, and Korbmacher C. Critical role of the mineralocorticoid receptor in aldosterone-dependent and aldosterone-independent regulation of ENaC in the distal nephron. American journal of physiology Renal physiology 321: F257–F268, 2021.

56. Pearce D, Manis AD, Nesterov V, and Korbmacher C. Regulation of distal tubule sodium transport: mechanisms and roles in homeostasis and pathophysiology. Pflugers Arch 474: 869–884, 2022.

57. Nesterov V, Dahlmann A, Krueger B, Bertog M, Loffing J, and Korbmacher C. Aldosterone-dependent and -independent regulation of the epithelial sodium channel (ENaC) in mouse distal nephron. American journal of physiology Renal physiology 303: F1289–1299, 2012.

58. Giacchetti G, Sechi LA, Griffin CA, Don BR, Mantero F, and Schambelan M. The tissue renin-angiotensin system in rats with fructose-induced hypertension: overexpression of type 1 angiotensin II receptor in adipose tissue. Journal of hypertension 18: 695–702, 2000.

59. Zenner ZP, Gordish KL, and Beierwaltes WH. Free radical scavenging reverses fructose-induced salt-sensitive hypertension. Integr Blood Press Control 11: 1-9, 2018.

60. Gordish KL, Kassem KM, Ortiz PA, and Beierwaltes WH. Moderate (20%) fructose-enriched diet stimulates salt-sensitive hypertension with increased salt retention and decreased renal nitric oxide. Physiological reports 5: 2017.

61. Masilamani S, Kim GH, Mitchell C, Wade JB, and Knepper MA. Aldosterone-mediated regulation of ENaC alpha, beta, and gamma subunit proteins in rat kidney [see comments]. Journal of Clinical Investigation 104: R19–R23, 1999.

62. Kleyman TR, Carattino MD, and Hughey RP. ENaC at the cutting edge: regulation of epithelial sodium channels by proteases. J Biol Chem 284: 20447–20451, 2009.

63. Passero CJ, Carattino MD, Kashlan OB, Myerburg MM, Hughey RP, and Kleyman TR. Defining an inhibitory domain in the gamma subunit of the epithelial sodium channel. American journal of physiology Renal physiology 299: F854–861, 2010.

64. Chen R, Sun W, Gu H, and Cheng Y. Aldosterone-induced expression of ENaC-alpha is associated with activity of p65/p50 in renal epithelial cells. J Nephrol 30: 73–79, 2017.

65. Gonzalez-Vicente A, Cabral PD, Hong NJ, Asirwatham J, Saez F, and Garvin JL. Fructose reabsorption by rat proximal tubules: role of Na(+)-linked cotransporters and the effect of dietary fructose. American journal of physiology Renal physiology 316: F473–F480, 2019.

66. Cabral PD, Hong NJ, Hye Khan MA, Ortiz PA, Beierwaltes WH, Imig JD, and Garvin JL. Fructose stimulates Na/H exchange activity and sensitizes the proximal tubule to angiotensin II. Hypertension 63: e68–73, 2014.

67. Ares GR, Kassem KM, and Ortiz PA. Fructose acutely stimulates NKCC2 activity in rat thick ascending limbs by increasing surface NKCC2 expression. American journal of physiology Renal physiology 316: F550–F557, 2019.

68. Komnenov D, and Rossi NF. Fructose-induced salt-sensitive blood pressure differentially affects sympathetically mediated aortic stiffness in male and female Sprague-Dawley rats. Physiological reports 11: e15687, 2023.

69. Rabie EM, Heeba GH, Abouzied MM, and Khalifa MM. Comparative effects of Aliskiren and Telmisartan in high fructose diet-induced metabolic syndrome in rats. Eur J Pharmacol 760: 145–153, 2015.

70. Kamari Y, Harari A, Shaish A, Peleg E, Sharabi Y, Harats D, and Grossman E. Effect of telmisartan, angiotensin II receptor antagonist, on metabolic profile in fructose-induced hypertensive, hyperinsulinemic, hyperlipidemic rats. Hypertens Res 31: 135–140, 2008.

71. Navarro-Cid J, Maeso R, Perez-Vizcaino F, Cachofeiro V, Ruilope LM, Tamargo J, and Lahera V. Effects of losartan on blood pressure, metabolic alterations, and vascular reactivity in the fructose-induced hypertensive rat. Hypertension 26: 1074–1078, 1995.

72. Soncrant T, Komnenov D, Beierwaltes WH, Chen H, Wu M, and Rossi NF. Bilateral renal cryodenervation decreases arterial pressure and improves insulin sensitivity in fructose-fed Sprague-Dawley rats. Am J Physiol Regul Integr Comp Physiol 315: R529–R538, 2018.

